# Transcriptional and epigenetic response of rapeseed (*Brassica napus* L.) to PEG-triggered osmotic stress

**DOI:** 10.1101/2024.08.19.608641

**Authors:** Melvin Prasad, Prateek Shetty, Avik Kumar Pal, Gábor Rigó, Kamal Kant, Laura Zsigmond, István Nagy, P. V. Shivaprasad, László Szabados

## Abstract

Drought hinders growth, development, and productivity of higher plants. While physiological and molecular background of plant responses to drought has extensively studied, the role of epigenetic modifications in response to dehydration remains largely elusive. In this study, we deciphered genome-wide transcriptomic and epigenetic responses of rapeseed (*Brassica napus* L.) to dehydration. High-throughput transcript profiling (RNA-seq) and chromatin immunoprecipitation followed by sequencing (ChIP-seq) of PEG-treated rapeseed plants revealed genome-scale changes in transcription and histone methylation patterns, focusing on histone H3 lysine 4 trimethylation (H3K4me3) and histone H3 tri-methylated lysine 27 (H3K27me3). We have identified large gene sets with altered transcript profiles and changed histone methylation marks in response to osmotic stress, revealed a close correlation between gain or loss of histone methylation and activation or repression of gene expression. Significant epigenetic regulation of Delta 1-Pyrroline-5-Carboxylate Synthetase (P5CS) genes, which control the key step in proline synthesis, was discovered as several PEG-induced *BnP5CSA* genes displayed enhanced H3K4me3 and/or H3K36me3 marks. Targeted bisulphite sequencing revealed that one *BnP5CSA* gene has stress-dependent gene body DNA methylation also. By integrating physiological, transcriptional and epigenetic data, our study facilitates better understanding drought response control in higher plants.

## Introduction

Plants are continuously exposed to changing environmental conditions and stresses that can negatively impact their growth and development. Abiotic stresses, including drought, salinity, and extreme temperatures, can have detrimental effects on plants, triggering reduced photosynthesis, cytotoxicity, disrupted essential metabolic and physiological processes, stunted growth, and nutrient-related disorders(He et al., 2018; Paes de Melo et al., 2022; Zhang et al., 2023). Drought, significantly affects plant growth, development, and yield. It leads to reduced biomass, poor seed quality, and lower crop productivity, posing a major threat to global food security(Iqbal et al., 2020; Kapoor et al., 2020). To survive and thrive under these conditions, plants have evolved various mechanisms to cope with drought stress by adjusting their physiological, biochemical, and genetic parameters(Takahashi et al., 2020). Epigenetic adjustments are reversible modifications affecting gene activation or silencing and represent significant regulatory aspect of plant adaptation to abiotic stress. In response to abiotic stresses, plants undergo various epigenetic modifications, such as changes in DNA methylation patterns, histone modifications, and the production of small RNAsClick or tap here to enter text. (Matzke and Mosher, 2014, Zhang et al., 2018, Chang et al., 2020, Miryeganeh, 2021).

DNA methylation controls gene expression, transposon silencing, and chromosome interactions. While DNA methylation in gene promoters typically represses transcription, in gene bodies it can enhance transcription(Zhang et al., 2006; Zhang et al., 2018; Lei et al., 2015; Williams et al., 2015; Lang et al., 2017). DNA methylation in gene bodies predominantly occurs in the CG context, and its function is not yet fully understood(Zhang et al., 2006; Takuno and Gaut, 2013; Lei et al., 2015; Zhang et al., 2018). DNA methylation changes enable plants to adapt to stress conditions faster by facilitating the expression of stress-responsive genes (Liu et al., 2010). Manipulating DNA methylation at specific loci may allow us to control gene expression and alter neighboring chromatin states.

Histone modifications, including acetylation, methylation, and phosphorylation, also play a crucial role in regulating gene expression in response to different environmental stimuli. These chemical modifications can alter chromatin compactness and gene accessibility, allowing the activation or repression of specific stress-responsive genes (Nunez-Vazquez et al., 2022). Histone acetylation on lysine residues in H3 or H4 decreases their affinity for DNA, reduces chromatin compactness, increases accessibility to transcription factors and subsequently tends to promote transcription (Shahbazian and Grunstein, 2007). Highly acetylated histones are often associated with active genes, while de-acetylated histones are more frequent in inactive genes. The effects of histone methylation are more complex, and depends on the amino acid residues involved. Three methylation of Lys4 in H3 (H3K4me3) is an active mark, allowing higher level of gene expression, while three methylation of Lys27 in H3 is considered a repressive mark, which appears frequently in repressed genes in heterochromatin(Nunez-Vazquez et al., 2022; Sun et al., 2022). Histone modifications are conserved and are mediated by similar enzymes in all eukaryotes: histone acetyltransferases (HATs) and histone deacetylases (HDACS) perform histone acetylation and deacetylation reactions, respectively, while histone methyltransferases (HMTs) aggregate and histone demethylases (HDMs) remove methyl groups. Activities of these enzymes are therefore important in determining histone modifications and can profoundly influence gene expression profiles in eukaryotic organisms(Pandey et al., 2002; Pontvianne et al., 2010; Nunez-Vazquez et al., 2022). The regulation of gene expression through epigenetic modifications is vital for plant adaptability to stress, as it enables a dynamic and reversible response to environmental changes(Abdulraheem et al., 2024). Reversible changes in H3K4me3 and H3K27me3 marks are often associated with changes in stress-dependent gene expression and were suggested to be essential components of adaptation to harmful conditions in higher plants(Kim et al., 2015; Nunez-Vazquez et al., 2022). Furthermore, abiotic stress can influence the production and function of small RNAs, such as microRNAs and small interfering RNAs, which target and regulate the expression of genes involved in stress (Matzke and Mosher, 2014). These small RNAs also mediate the silencing of transposable elements, thereby protecting the genome from stress-induced instability (Sunkar et al., 2007; Kamthan et al., 2015).

Epigenetic changes induced by abiotic stress can sometimes be inherited to subsequent plant generations, a phenomenon known as transgenerational epigenetic inheritance(Hauser et al., 2011; Godwin and Farrona, 2020). This inheritance can help subsequent generation adapt more effectively to similar stress conditions, thereby providing a survival advantage. Characterizing these epigenetic adjustments is essential for understanding plant responses to abiotic stress and revealing the molecular basis of plant adaptation to extreme environmental conditions (Ashapkin et al., 2020; Rai et al., 2023). Engineering epigenetic variation to develop improved abiotic stress-tolerant crop varieties is a promising approach which can contribute to long-term plant resilience during global climatic changes (Singroha and Sharma, 2019; Kakoulidou et al., 2021).

A significant amount of our understanding of epigenetic regulation has been derived from studies on model organisms like Arabidopsis. Knowledge of the epigenetic profiles of crops under stress can facilitate the development of novel strategies for creating stress-tolerant varieties through epigenetic engineering or targeted breeding programs. Polyploidy-dependent changes in epigenetic regulation such as hypomethylation can promote responses to adverse environmental stimuli(Yuan et al., 2020; Wang et al., 2021). Advancements in high-throughput sequencing techniques have revolutionized the identification of epigenetic changes on genomic level and facilitate our understanding of their impact on gene expression regulation(McCombie et al., 2019). Information on stress-related epigenetic regulation of crops with large and polyploid genomes such as the allotetraploid rapeseed (*Brassica napus*) is however scarce. Comprehensive epigenome map of rapeseed has been published, providing genome-scale information on histone modifications, DNA methylation and RNA polymerase II occupancy. Assimetric histone modifications have been found between the rapeseed An and Cn subgenomes, as well as between various tissue types (Zhang et al., 2021). Differences in epigenetic marks between rapeseed subgenomes have recently been reported which correlated with changes in transcript profiles (Hu et al., 2023). Circadian oscillation of histone modifications in rapeseed have also been described, pointing to differential regulation of homologous gene pairs of the subgenomes(Xue et al., 2023). Another study characterized the epigenetic modification of transposable elements in the evolution of the allopolyploid nature of rapeseed (Xiao et al., 2022). While these studies have generated valuable datasets and revealed important information about the epigenome map of rapeseed, and revealed information on chromatin states and regulatory elements on genomic scale, epigenetic control on responses to environmental stresses such as drought in rapeseed is still unknown.

In this study, the widely cultivated rapeseed cultivar Westar (Klassen et al., 1987) was subjected to osmotic stress using PEG treatment. Comprehensive transcriptome analyses identified drought-responsive differentially expressed genes (DEGs) involved in tolerance mechanisms. We performed H3K4me3 and H3K27me3 ChIP-seq assays on leaf tissue to characterize the subgenomic distribution of these marks in control and PEG-treated plants. This allowed us to link the enrichment or depletion of active and repressive histone modifications to their target genes, thereby regulating differential gene expression. Our RNA-seq and ChIP profiling revealed that several rapeseed P5CSA genes, controlling proline accumulation, are under epigenetic control. Besides histone modifications, rapid redistribution of DNA methylation marks upon osmotic stress in one of the P5CSA genes could be confirmed. This study represents the first transcriptomic and genome-wide H3K4me3 and H3K27me3 ChIP-seq analysis in rapeseed under osmotic stress, broadening our understanding of epigenetic marks in plant stress responses. Our findings provide valuable insights into the genetic and molecular mechanisms of stress response of an allotetraploid crop with complex genome, and offer genetic resources for the improvement of high-yielding, drought-tolerant oilseed rapeseed varieties.

## Materials and methods

### Plant material and growth conditions

Plants of *Brassica napus* var. Westar (Klassen et al., 1987) were grown on hydrophonic system using Hoagland nutrient solution as described(Wang et al., 2019). Plants were maintained and all treatments were performed in a growth chamber (Fytoscope SW, PSI, Czech Republic), under precisely controlled conditions. Illumination was provided with LED panels with photon flux of 160 μmol of photons m-2s-1 and a short-day light cycle (8 h light at 22°C / 16 h darkness at 20°C). Four-weeks-old plants were subjected to osmotic stress by replacing the standard Hoagland nutrient solution with solution supplemented by 20% PEG8000. Plants were allowed to recover from osmotic stress by replacing the PEG-containing solution with standard Hoagland solution. Leaf samples of PEG-treated and control plants were collected at various time points, specifically at 6 hours, 24 hours, 48 hours, 3 days, 5 days, 7 days, 10 days, and 15 days, immediately frozen in liquid nitrogen and stored until processing (workflow is show in Figure S1A). Experiments were repeated three times (biological replicates).

### Determination of proline content

Proline content was determined as described with minor modifications(Kovács et al., 2019). 50 mg of leaf tissue was ground with liquid nitrogen, 1 ml of 1% sulfosalicylic acid was added and mixed by vortex. The mixture was centrifuged at 13000 rpm for 10 min at 4°C, and 200 μl supernatant was mixed with 400 μl of 1.25 % ninhydrin reagent (ninhydrin dissolved in 80 % acetic acid). The samples were incubated in the dry block heater bath at 95°C for 30 min, and immediately placed on ice for several minutes. Proline content was determined by measuring the absorbance of the reaction product at 520 nm using Thermo Scientific, Multiscan Go Microplate Spectrophotometer. To draw the standard curve of proline, concentrations of 0.5, 0.25, 0.125, 0.025, 0.03125 and 0 mM proline were used as reference. The content of proline was measured with five technical replicates.

### Determination of malondialdehyde

Lipid peroxidation rates were measured by the thiobarbituric acid-reactive substances (TBARS) assay as described (Heath and Packer, 1968). 100 mg leaf tissue was homogenized in 1 ml of 0.1% trichloroacetic acid (TCA) containing 0.4% butylhydroxytoluene, centrifuged at 13,000 rpm for 20 minutes. 1 ml of 20% TCA containing 0.5% thiobarbituric acid (TBA) was added to 250 μl supernatant, mixed and incubated at 96 °C for 30 min. Absorbance was measured at 532 nm using a Multiskan GO microplate reader (Thermo Fisher Scientific) and calculated by subtracting the nonspecific absorbance measured at 600 nm. Malondialdehyde (MDA) concentration was calculated using the extinction coefficient ε532 = 155 mM−1 cm−1. Three biological replicates were made.

### Determination of photosynthetic parameters

Chlorophyll fluorescence measurements on light-adapted leaves were carried out directly in the growth chamber using a portable MultispeQ v1 device (PhotosynQ) controlled by the PhotosynQ platform software (Kuhlgert et al., 2016), using a red AL (200 μmol photons m−2 s−1) and a 500-ms saturation pulse (3000 μmol photons m−2 s−1).

### RNA Extraction and RT-qPCR Analysis

Total RNA was isolated using a GeneJET RNA Purification Kit (Thermo scientific) following the manufacturer’s protocol. One μg of DNase-treated RNA was used for cDNA synthesis using the High-Capacity cDNA Reverse Transcription Kit (Applied Biosystems). qRT-PCR was performed on 3 μl of 20x diluted cDNA templates using 5x HOT FIREPol EvaGreen qPCR Mix Plus (Solis Biodyne) in a final volume of 10 μl, and Bio-Rad CFX96 Touch Deep Well Real-Time PCR System. Mean values of Actin and GAPDH Ct were used as internal reference. Normalized relative transcript levels were determined using the 2^-ΔΔCt^ method (Livak and Schmittgen, 2001). Experiments were repeated with three biological replicates. Oligonucleotides used in this study are listed in **Table S1**.

### RNA-seq analysis

Total RNA was isolated using a GeneJET RNA Purification Kit (Thermo scientific) following the manufacturer’s protocol. Approximately 3μg of total RNA from each sample was subjected to the RiboMinus Eukaryote Kit (Qiagen, Hilden, Germany) to remove ribosomal RNA prior to the construction of the RNA-seq libraries. RNA-seq libraries were prepared using an RNA-seq Library Prep Kit for Illumina (New England Biolabs). The libraries were sequenced on an Illumina HiSeq 4000 platform using a SE150-bp read module. The raw sequencing reads were filtered and trimmed using the default parameters of Fastp (version 0.23.2) (https://academic.oup.com/bioinformatics/article/34/17/i884/5093234). The filtered clean data were then pseudo-aligned against the transcript models for *Brassica napus* var. Westar downloaded from (http://cbi.hzau.edu.cn/cgi-bin/rape/download_ext) using Kallisto (version 0.50.1). Differentially expressed genes (DEGs) were identified using DESeq2 (version 1.42.1 (http://bioconductor.org/packages/release/bioc/html/DESeq.html) with the criteria of q value (adjusted p value, Benjamini-Hochberg method) <0.05 and |log_2_ Fold Change (FC)| ≥2)(Love et al., l., 2014). Functions of the DEGs were investigated with Gene Ontology (GO) and Kyoto Encyclopedia of Genes and Genomes (KEGG) pathway enrichment analysis using topGO (version 4.4) and ClusterProfiler (version 4.12.2)(Alexa and Rahnenfuhrer, 2010; Wu et al. 2021), respectively. Significant GO terms and KEGG pathways were identified with the criterion of q value <0.05. Sequence data were deposited in the National Center for Biotechnology Information (NCBI) SRA database (Accession number: PRJNA1144338)

### ChIP-seq and ChIP-qRT PCR

ChIP assays were performed as previously described by Song et al. (2016) with minor modifications. In brief, 2 to 3 g of 30-day old plant leaves (Mock and PEG treated) were crosslinked with 1% formaldehyde for 20 min under vacuum, quenched with freshly prepared 2M glycine and ground into fine powder in liquid nitrogen. Chromatin was isolated and sheared into 200 to 500 bp DNA fragments by sonication. The sonicated chromatin was immunoprecipitated with one of the following antibodies (5 μg): Anti-trimethyl-Histone H3 (Lys4) (Merck, 07-473), Anti-Histone H3K27me3 (Active motif, 39155), Anti-Histone H3 (di methyl K9) (Abcam, ab1220), Anti-Histone H3 (tri methyl K36) (Abcam ab9050), Anti-Histone H4 (acetyl K5) (Abcam, ab51997) or Anti-Myc tag antibody (used as IgG negative control, Abcam, ab9E11) and with 25 μl of Dynabeads Protein G (Invitrogen, 10003D) for 12 h at 4 °C with rotation. The precipitated chromatin DNA was then purified by phenol– chloroform-isoamyl alcohol extraction and recovered by ethanol precipitation. The ChIP DNA was prepared for sequencing or qPCR. Two biological replicates were used for ChIP-seq, and three biological replicates were used for ChIP–qPCR. ChIP-IgG was used for normalising the values. the primers used for ChIP qPCR are listed in **Table S2**. Actin (BnaC09T0320800WE) was used as a negative control.

### ChIP-seq data analysis

At least 10 ng of ChIP DNA was used for library preparation. ChIP sequencing libraries were constructed using NEBNext Ultra II DNA Library Prep Kit (New England Biolabs Inc, E7103) following the manufacturer’s instructions. Constructed libraries were sequenced using Illumina NovaSeq 6000 and paired-end reads were obtained at Geninus Bowtie2 software (https://www.kr-geninus.com) (Langmead and Salzberg, 2012) and were used to align the sequencing reads of ChIP-seq to the *Brassica_napus* reference genome (http://cbi.hzau.edu.cn/rape/download_ext/westar.genome.fa) using default parameters. The peak in different conditions and differentially changed peaks were called by MACS software (Zhang et al., 2008). The nomodel parameter was set, and the d-value parameter was set at 200. The resulting wiggle files, which represent counts of ChIP-Seq reads across the reference genome, were normalized for sequencing depth by dividing the read counts in each bin by the millions of mapped reads in each sample and were visualized in the IGV genome browser. The Diffbind was used to Compute differentially bound sites between PEG-treated and control conditions from multiple ChIP-seq experiments using affinity (quantitative) data. For the analysis of histone modification profiles between the 24-hour PEG-treated samples and the mock-treated samples, the Galaxy platform was employed (<u>usegalaxy.eu</u>). Sequence data were deposited in the National Center for Biotechnology Information (NCBI) SRA database (Accession number: PRJNA1144218).

### Targetted DNA methylation

Primers for the targeted DNA methylation were designed using Bisulphite-primer-seeker tool (https://zymoresearch.eu). DNA from the rapeseed was isolated using DNA extraction kit Nucleon PhytoPure (Merck). Bisulphite conversion of DNA was carried using the kit EZ DNA Methylation-Gold Kit (Zymoresearch, D5005). Bisulphite-converted DNA was used as a template for the targeted DNA amplification. PCR fragments were purified and used for sequencing. Adapters were removed from the deep sequenced PCR products by cutadapt tool (Martin, 2014). For alignment the reads, the target sequence was converted by Bismark aligner. Adapter trimmed sequences (90-100 bp) were aligned to target site using Bismark aligner tool with default parameters (Krueger and Andrews, 2011). DNA methylation status of the targeted site was extracted and coverage reports were generated using the Bismark aligner tool. The obtained results were analysed using methylation package ViewBS (Huang et al., 2018). Sequence data were deposited in the National Center for Biotechnology Information (NCBI) SRA database (Accession number: PRJNA1144360).

### Statistical analyses

Statistical analysis was performed using two-way analysis of variance (ANOVA) followed by the post hoc Tukey HSD test (P <0.05) using the Graphpad prism software (version 8). The data presented in graphs represent the mean value ± standard error of three independent experiments.

## Results

### 1. Consequences of PEG-induced osmotic stress on rapeseed

Water deprivation during drought generates osmotic stress in plants can be simulated by Polyethylene glycol (PEG) treatment. Additionally, PEG provides consistent, reproducible, homogeneous stress conditions (Wang et al. 2019). To investigate the impact of osmotic stress on Westar cultivar of rapeseed, a hydroponic system was employed with 20 % PEG8000 as osmotic agent. Samples for physiological and molecular assays were collected at diverse timepoints (**Figure S1A**). Rapeseed plants exhibited immediate signs of water loss, which manifested as noticeable wilting within several hours of PEG treatment (**Figure 1A**).

**Figure 1:**
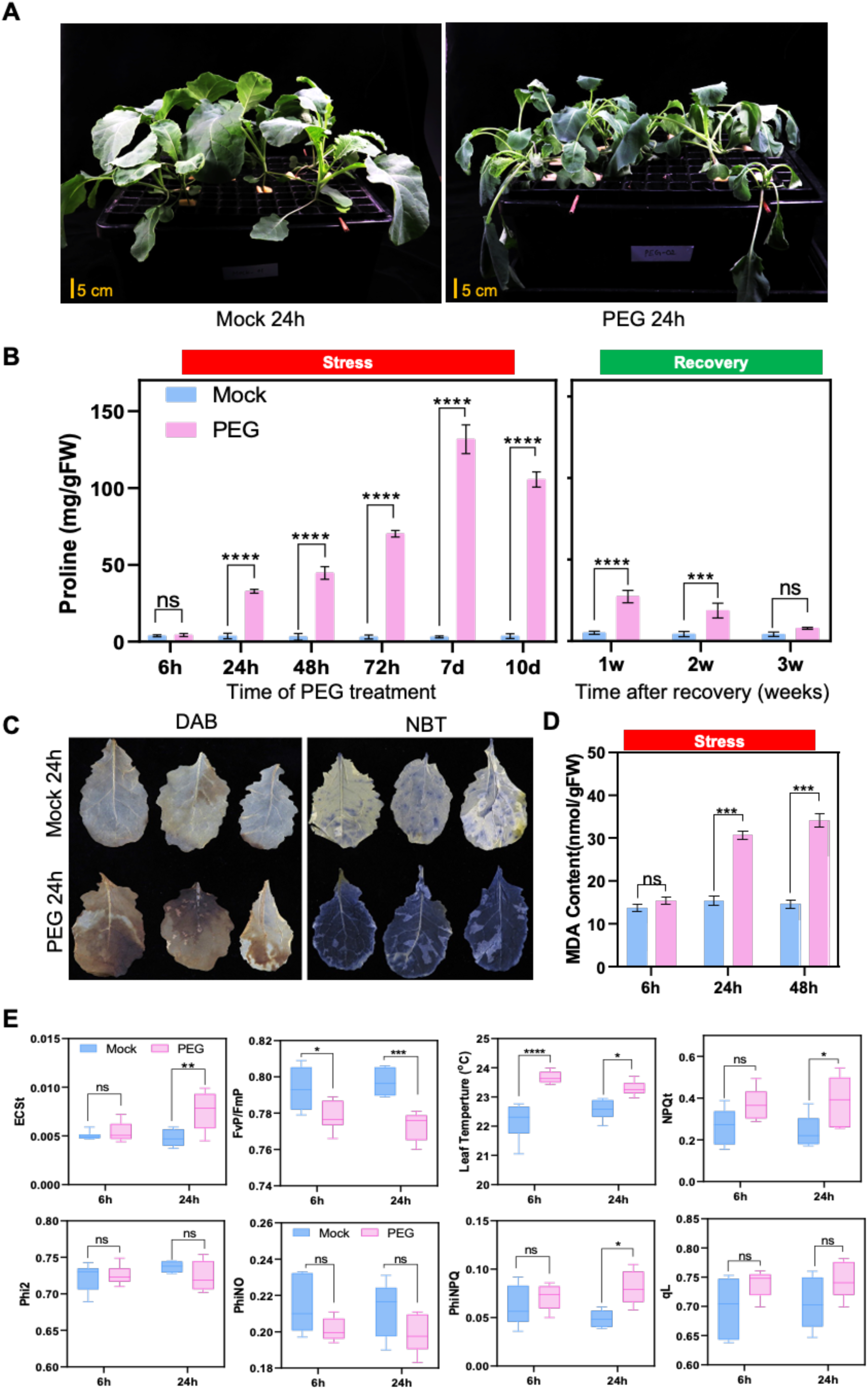
Effect of Osmotic Stress on Rapeseed Plants. **A)** Morphological changes in rapeseed plants after 24 hours of 20% PEG treatment in hydroponic culture. Note the differences in leaf appearance, with leaves losing turgor as a consequence of osmotic stress, **B)** Proline levels in leaves of rapeseed plants during osmotic stress and after recovery. Left panel shows changes in proline content up to 10 days of PEG traement. Right panel shows proline content after 1-2-3 weeks of recovery. **C)** Accumulation of Reactive Oxygen Species (ROS) in stressed and control rapeseed plants (Mock). Histochemical detection of hydrogen peroxide using 3,3-diaminobenzidin (DAB) staining and superoxide levels using nitroblue tetrazolium (NBT) staining in rapeseed plants. **D)** Lipid peroxidation measured as Malondialdehyde (MDA) accumulation in leaves of rapeseed plants subjected to 6, 24 and 48 hours of stress, compared to controls. **E)** Effect of osmotic stress on photosynthetic parameters of PSII in rapeseed plants: energy-dependent quenching (ECSt), maximum quantum efficiency of PSII (FvP/FmP), quantum yield of non-regulated energy dissipation (PhiNO), quantum yield of PSII photochemistry (Phi2), leaf temperature (C°), non-photochemical quenching (NPQ), photochemical quenching coefficient (qL), quantum yield of regulated energy dissipation (PhiNPQ). Each data point represents the average of six replicates. Bars represent the mean ± SE of three independent experiments. Statistical analysis was performed using a two-way ANOVA followed by Tukey’s multiple comparison test. Columns labeled with "ns" indicate no significant difference, while asterisks (*) denotes p < 0.05, (**) p < 0.01, (***) p < 0.001, and (****) p < 0.0001 indicate statistically significant differences among treatments.

Proline accumulation in higher plants is a characteristic response to extreme conditions such as drought or high soil salinity and is often considered as marker for altered physiological homeostasis (Alvarez et al., 2022). PEG-treated plants displayed a significant increase in proline content compared to their respective control counterparts. Leaf samples had the highest level of proline accumulation in response to PEG treatment, while stems and roots had negligible proline (**Figure S1B**), suggesting that metabolic consequences of the osmotic stress was strongest in leaves. For this reason subsequent assays were performed with leaves. When time-dependent proline accumulation was tested, a significant rise in proline content was particularly noticeable in the early stages of stress exposure, with a gradual and sustained increase observed from the 6-hour time point through to the 7-day interval. Following this peak, proline levels declined, which became more apparent after 10 days of PEG treatment. When PEG was replaced with standard nutrient solution, proline levels were reduced to nearly control in three weeks (**Figure 1B**).

Accumulation of reactive oxygen species (ROS) in stressed plants is known to generate oxidative damage, which was monitored in PEG-treated rapeseed. Hydrogen peroxide and superoxide levels were analysed with DAB and NBT histochemical reactions. Both assays generated intensive color reactions in PEG-treated rapeseed, but not in control plants, indicating considerable ROS accumulation as result of osmotic stress (**Figure 1C**). Additionally, We observed a significantly higher MDA content in the PEG-treated plants at 24h and 48h of stress exposure compared to the control group, indicating extensive oxidative damage (**Figure 1D**).

To further characterize stress-dependent physiological changes, photosynthetic parameters were compared in PEG-treated and control rapeseed plants. Significant changes were observed in various parameters under osmotic stress comparing to controls. There was a notable decrease in maximum quantum efficiency of PSII in the light-adapted state (FvP/FmP) and quantum yield of non-regulated energy dissipation (PhiNO) in the stressed plants. Electron transport rate through PSII (ECSt), non-photochemical quenching (NPQ), and quantum yield of regulated energy dissipation (PhiNPQ) as well as leaf temperatures were higher in PEG-treated plants. No significant differences were detected in photochemical quenching coefficient (qL) and quantum yield of PSII photochemistry (Phi2) (**Figure 1E**). These findings suggest that PEG influences specific changes in the photosynthetic performance of rapeseed plants, particularly in aspects related to energy dissipation and heat regulation, while maintaining steady-state photochemical efficiency.

### PEG -induced transcriptomic changes

Proline levels are determined by the activities of both biosynthetic and catabolic pathways, controlled by delta-1-pyrroline carboxylate-5-synthase (P5CS), and by proline dehydrogenase (PDH) enzymes, respectively (Szabados and Savouré, 2010; Mattioli et al., 2022). In Arabidopsis, P5CS is encoded by two genes, *P5CS1* and *P5CS2* with remarkable differences in their expression regulation: while *P5CS1* is induced by high osmolarity and ABA, *P5CS2* expression is less influenced by environmental signals (Savouré et al., 1997; Strizhov et al., 1997; Kovács et al., 2019). In contrast to Arabidopsis, rapeseed possesses 10 P5CS genes, which belong to the P5CSA and P5CSB subgroups (**Table S6**). Transcript levels of the A and B-types of rapeseed P5CS has been determined by RT-QPCR, although this method is unsuitable for distinguishing between individual genes in these subfamilies due to their high sequence identities. Both *BnP5CSA* and *BnP5CSB* gene groups were rapidly activated in PEG-treated plants, with their transcript levels remaining elevated throughout the high osmotic conditions (**Figure 2**). In *Arabidopsis*, stress-dependent *P5CS1* induction relies on both ABA-dependent and independent signals (Savouré et al., 1997). To investigate whether similar regulatory mechanisms are activated in rapeseed during osmotic stress, the expression of selected ABA and stress-responsive genes was monitored in PEG-treated and control plants.

**Figure 2.**
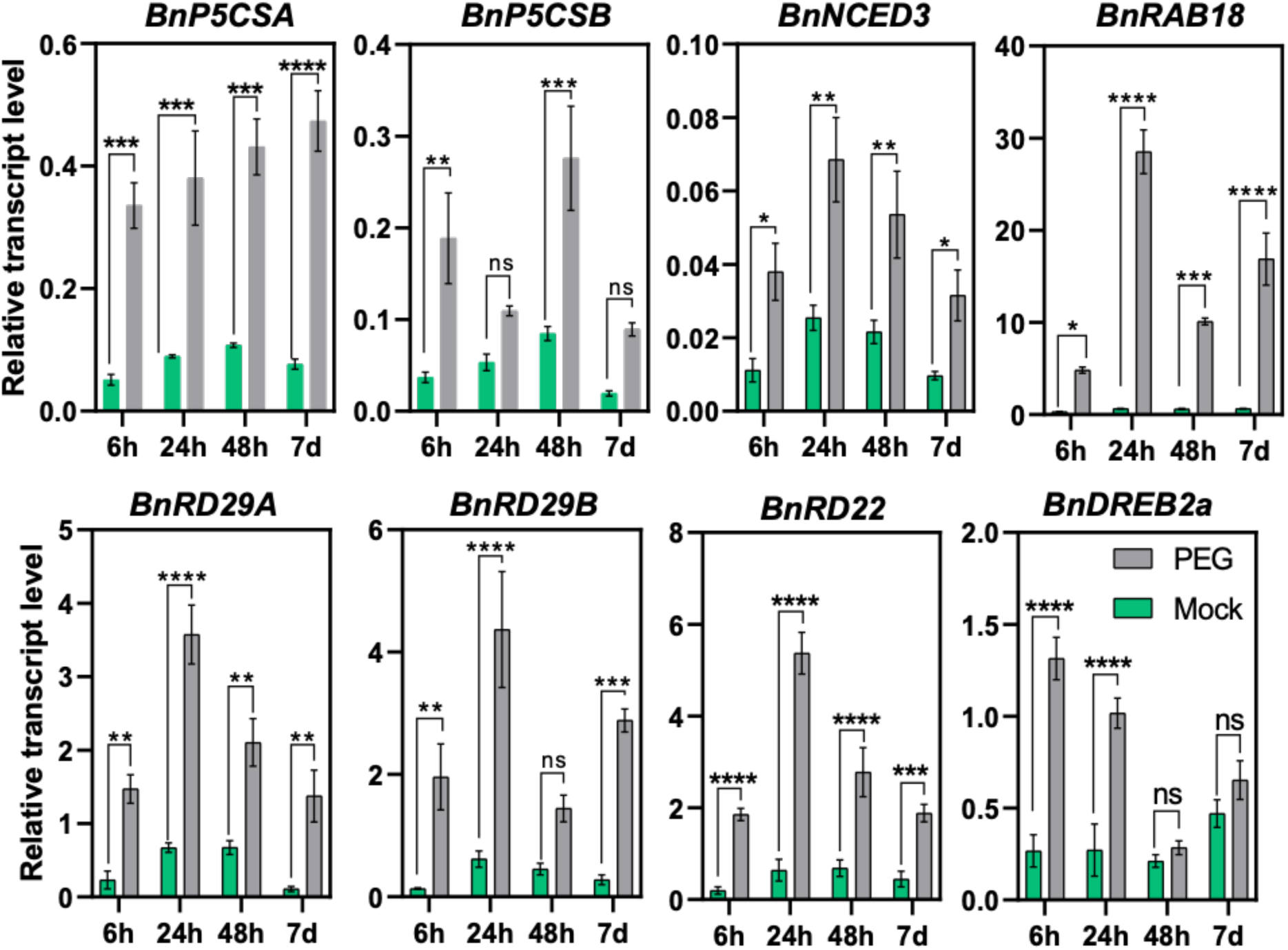
Response of Stress Marker Genes to PEG-Induced Osmotic Stress. Transcript levels of selected genes were determined by qRT-PCR in PEG-treated and control rapeseed plants: *BnP5CSA, BnP5CSB, BnNCED3, BnRAB18, BnRD29A, BnRD29B, BnRD22,* and *BnDREB2a*. Data represent the fold induction of each gene by osmotic stress relative to the *BnACT* transcript levels as a reference. Bars represent the mean ± SE of three independent experiments. Statistical analysis was performed using a two-way ANOVA followed by Tukey’s multiple comparison test. Columns labeled with "ns" indicate no significant difference, while asterisks (*) denotes p < 0.05, (**) p < 0.01, (***) p < 0.001, and (****) p < 0.0001 indicate statistically significant differences among treatments.

In Arabidopsis, ABA biosynthesis is regulated by the stress-responsive *NCED3* (Finkelstein, 2013). Transcript levels of *BnNCED3* were enhanced by PEG throughout the treatment, suggesting activation of ABA biosynthesis in rapeseed leading to enhanced ABA-dependent transcription. Arabidopsis genes *RD29A, RD29B, RD22,* and *RAB18* are well-characterized drought and ABA-induced genes (Takahashi et al., 2018). In rapeseed, *BnRD29A, BnRD29B, BnRD22,* and *BnRAB18* were quickly induced by PEG, with their transcript levels remaining elevated under high osmotic conditions. In Arabidopsis, *DREB2 TFs* are induced by water deprivation but not by ABA and control drought-dependent regulons (Liu et al., 1998). Rapeseed *BnDREB2A* was induced after 6 and 24 hours of PEG treatment, indicating an active ABA-independent regulation under high osmotic conditions (**Figure 2**). These transcript data provide compelling evidence that the PEG treatment generates substantial changes in gene expression patterns and is suitable for conducting transcriptomic and epigenetic investigations in rapeseed.

### 2. Genome-wide transcript analysis of rapeseed

To get information on genome-wide expression changes of rapeseed during osmotic stress, RNA-seq analysis was performed on four-weeks-old plants subjected to 6-hour and 24-hour PEG treatments and their respective controls. The RNA-seq quality control (QC) reports, detailing the counts of raw reads, trimmed reads, and mapping quality, are summarized in **Table S3**. Principal Component Analysis (PCA) of the RNA-seq data demonstrated that replicates cluster together and were distinctly separated from their respective treatment groups (**Figure 3A**). Genes with differential expression patterns could be clustered together as displayed by a heatmap (**Figure 3B).** For the 6-hour PEG-treated samples, our analysis identified 5473 Differentially Expressed Genes (DEGs), employing a statistical significance threshold of pAdj: 0.05 and a fold change cutoff of 2.0. In response to PEG treatment, 2798 genes were upregulated, while 2675 genes were downregulated (**Figure 3C, Figure S2A**). A comprehensive list of the genes displaying differential expression at 6 hours post-PEG treatment can be found in the **Supplementary datasets**. Gene ontology (GO) analysis was conducted to discern the functional attributes of the DEGs. Six hours of PEG treatment lead to significant upregulation of genes associated with transcriptional activity, DNA binding, chromatin binding, and response to various stress and stimuli (**Figure S2B**). Genes linked to transferase activity, protein kinases, catalytic processes, and metabolic pathways were downregulated by 6 h PEG treatment (**Figure S2C**). We conducted KEGG analysis on the DEGs identified after 6 hours of PEG treatment, revealing significant changes in the expression of genes associated with various metabolic and signal transduction pathways (**Figure S3**). Transcriptomic analysis of the 24-hour samples revealed 3379 DEGs, with 1179 genes being upregulated and 2200 genes downregulated (**Figure 3D, Figure S2A**). Gene ontology analysis of the 24-hour DEGs identified an enrichment of genes responsible for transcriptional activity, DNA binding, and stress response (**Figure S4A**), while there was a reduction in the expression of genes associated with transcription regulation, DNA binding, transport processes, and lipid metabolic pathways (**Figure S4B**). KEGG analysis revealed alterations in the genes involved in metabolic and signal transduction pathways, consistent with those observed at the 6-hour treatment (**Figure S5**).

**Figure 3.**
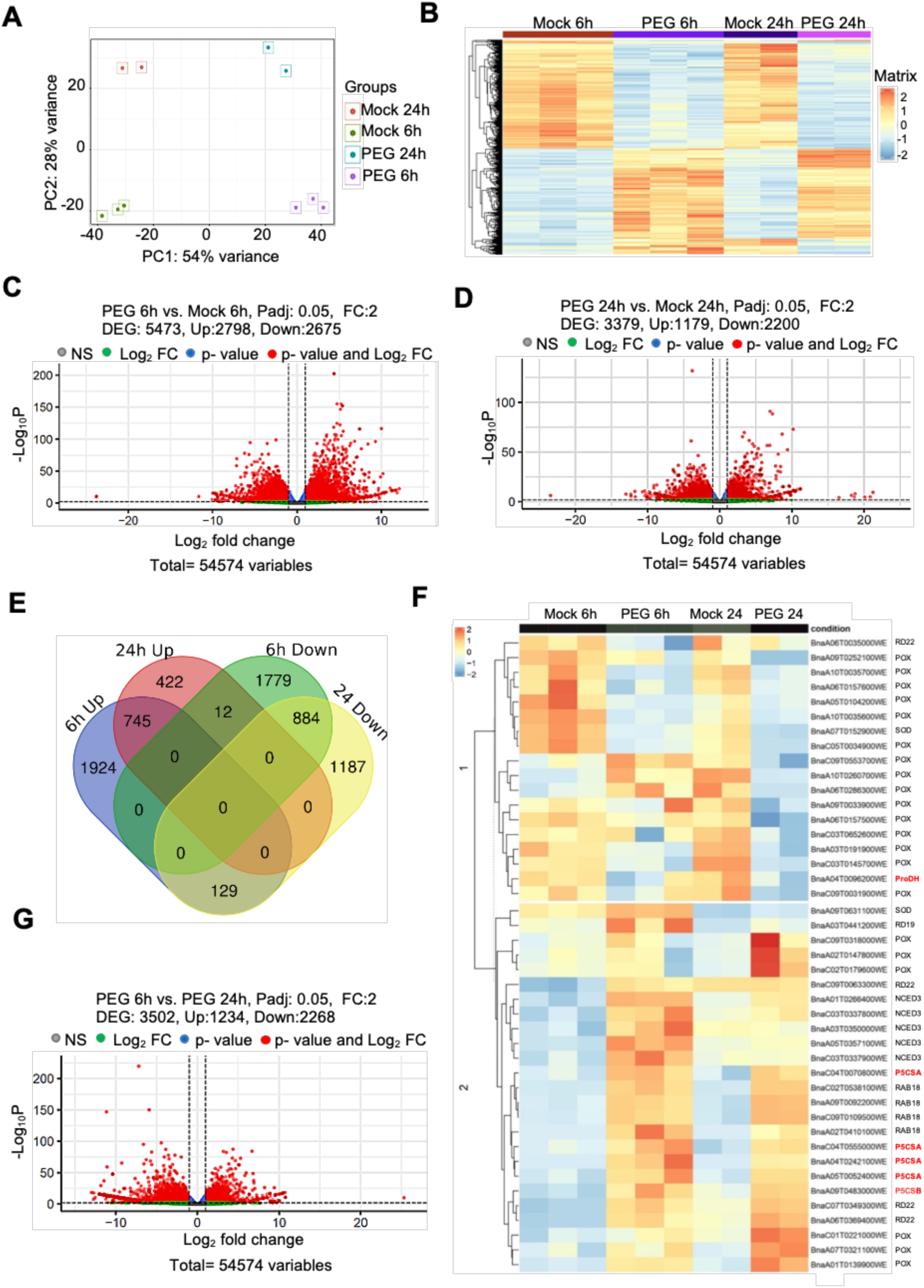
RNA-Seq Analysis of Rapeseed in Response to Osmotic Stress. **A)** Principal Component Analysis (PCA) of transcript datasets obtained from Mock and PEG-treated plants (6h and 24h). PCA revealed distinct expression signatures for each condition. The percentage of variation is indicated on the PC1 and PC2 axes. **B)** Heatmap representation of gene expression, calculated as the average Reads Per Kilobase of transcript per Million reads mapped (RPKM) value from the biological replicates. **C, D)** Volcano plot Analysis of differentially expressed genes (DEGs) in Mock and PEG-treated samples after 6h (C) and 24h (D) of PEG treatments. DEGs were determined by DESeq2, plots were generated with the R package ggplot2, display downregulated genes in blue and upregulated genes in red (p adj <0.05, fold change (fc) > 2). **E)** Venn diagram of DEGs illustrating the overlap and unique sets of genes regulated at 6h and 24h of PEG treatments. **F)** Expression heatmap of rapeseed stress-responsive genes at 6h and 24h after PEG treatment. **G)** Volcano Plot comparison of DEGs between 6h and 24h in Mock and PEG-treated samples (p adj <0.05, fold change (fc) > 2). The plot depicts down and upregulated genes in blue and red, respectively.

A comparative analysis between the 6-hour and 24-hour samples revealed that 745 genes were upregulated at both time points in response to the stress treatment, indicating a sustained stress response. Similarly, 884 genes displayed downregulation after 6-hour and 24-hour PEG, suggesting a conserved negative transcriptional response of these genes to osmotic stress. Intriguingly, a subset of 129 genes exhibited upregulation after 6-hour PEG treatment, which became downregulated after 24-hour of PEG (**Figure 3E**). Gene Ontology (GO) enrichment analyses showed, that genes with transcription regulatory, DNA binding, response to stress, response to stimuli, lipid transport and developmental and metabolic process were overrepresented among the upregulated genes (**Figure S6A**). For the genes downregulated after 6-hour and 24-hour PEG treatment, we identified those involved in metal ion transport and catalytic activity (**Figure S6B**). KEGG analysis showed that these genes are primarily involved in plant hormone and signal transduction pathways **(Figure S7A-B).** The genes, which were upregulated at 6 hours and downregulated at 24 hours, are mainly involved in transcription regulation, DNA binding, the ubiquitin ligase complex, as well as signal transduction pathways (**Figure S8A-B**). Twelve genes were initially downregulated at the 6-hour time point which experienced upregulation at the 24-hour of PEG treatment **(Figure 3E)**.

In order to get information on regulatory genes implicated in gene expression control, we outlined the differential expression patterns of crucial defense-related transcription factors and chromatin remodeler genes in response to PEG treatment. Transcripts associated with MYB, NAC, DREB, and HD-ZIP-type factors could exhibit either upregulation or downregulation. Notably, a substantial portion of transcripts from the MYC and ABF/ABI gene families displayed upregulation at both the 6-hour and 24-hour time points following PEG treatment. In contrast, a majority of WRKY transcripts exhibited downregulation (**Figure S9**). Furthermore, we assessed the expression of genes implicated in chromatin modifications. Transcript levels of several HDAC, HAC1, DRM2, JMJ, SUVH6, ATX, and SUVR genes showed higher expression, whereas CLASSY1-like, MET1, DME, CMT2, HAC1, and HDAC1 genes displayed reduced expression under PEG stress (**Figure S10**).

The expression patterns of a subset of drought-responsive defense-related genes were analyzed at 6h and 24h under drought stress. A heatmap presented the temporal and spatial expression patterns of 43 differentially expressed genes (DEGs). Most transcripts for P5CSA, NCED3, RD22, RD19, RAB18, and SOD exhibited a positive induction in response to PEG-induced stress. However, most transcripts for POX and ProDH were downregulated under these conditions (**Figure 3F**).

We further compared the gene sets differentially misregulated in 6 h and 24 h PEG-treated plants. We identified 3502 DEGs, with 1234 genes displaying upregulation and 2268 genes exhibiting downregulation after 24 hours of PEG when compared to 6-hour treatments **(Figure 3G, Figure S11A)**. GO analysis of genes upregulated in 24 h comparing 6 h samples revealed differences in enrichment in categories associated with transcription regulation, chromatin binding, catalytic and peptidase activities. GO terms which were identified in genes which were downregulated in 24 h PEG comparing to 6 h PEG were transcription regulators, DNA binding, oxidoreductase and catalytic activities (**Figure S11B, C**). These data revealed substantial changes in gene expression regulation of rapeseed plants exposed to short and long-term osmotic stress.

### 4. Epigenetic Profiling of H3K4me3 and H3K27me3 Marks in Response to PEG treatement

Histone modifications play pivotal roles in the regulation of gene expression governing various aspects of plant physiology, growth, development, and defense mechanisms. To identify genome-wide changes in epigenetic features of rapeseed associated with osmotic stress, ChIP-seq analysis was performed on leaf samples derived from plants subjected to 24 h PEG treatment and their respective controls. Comprehensive genome-wide profiling of two key histone marks was performed identifying histone H3 lysine 4 trimethylation (H3K4me3) and histone H3 lysine 27 trimethylation (H3K27me3), associated with gene activation and gene repression, respectively. The ChIP-seq data encompassed a total of 10 libraries, which comprised 8 for immunoprecipitated DNA and 2 for input DNA. These libraries collectively yielded approximately 212 million paired-end reads, with an average of roughly 15-21 million reads per sample (**Table S4**). Notably, an average of 76% of the reads in each library could be successfully mapped to the Rapeseed reference genome (BnPIR: http://cbi.hzau.edu.cn/bnapus). Comparing the histone modification profiles of the 24-hour PEG-treated and mock-treated samples revealed distinct variations in the principal components (PCA), effectively differentiating between the two histone modifications **(Figure 4A)**. Furthermore, the replicates displayed a consistent signal for H3K4me3 and H3K27me3, indicating robust experimental reproducibility. The pairwise correlation among the demultiplexed samples exhibited high values, ranging from 0.71 to 1.0 **(Figure 4B)**.

**Figure 4:**
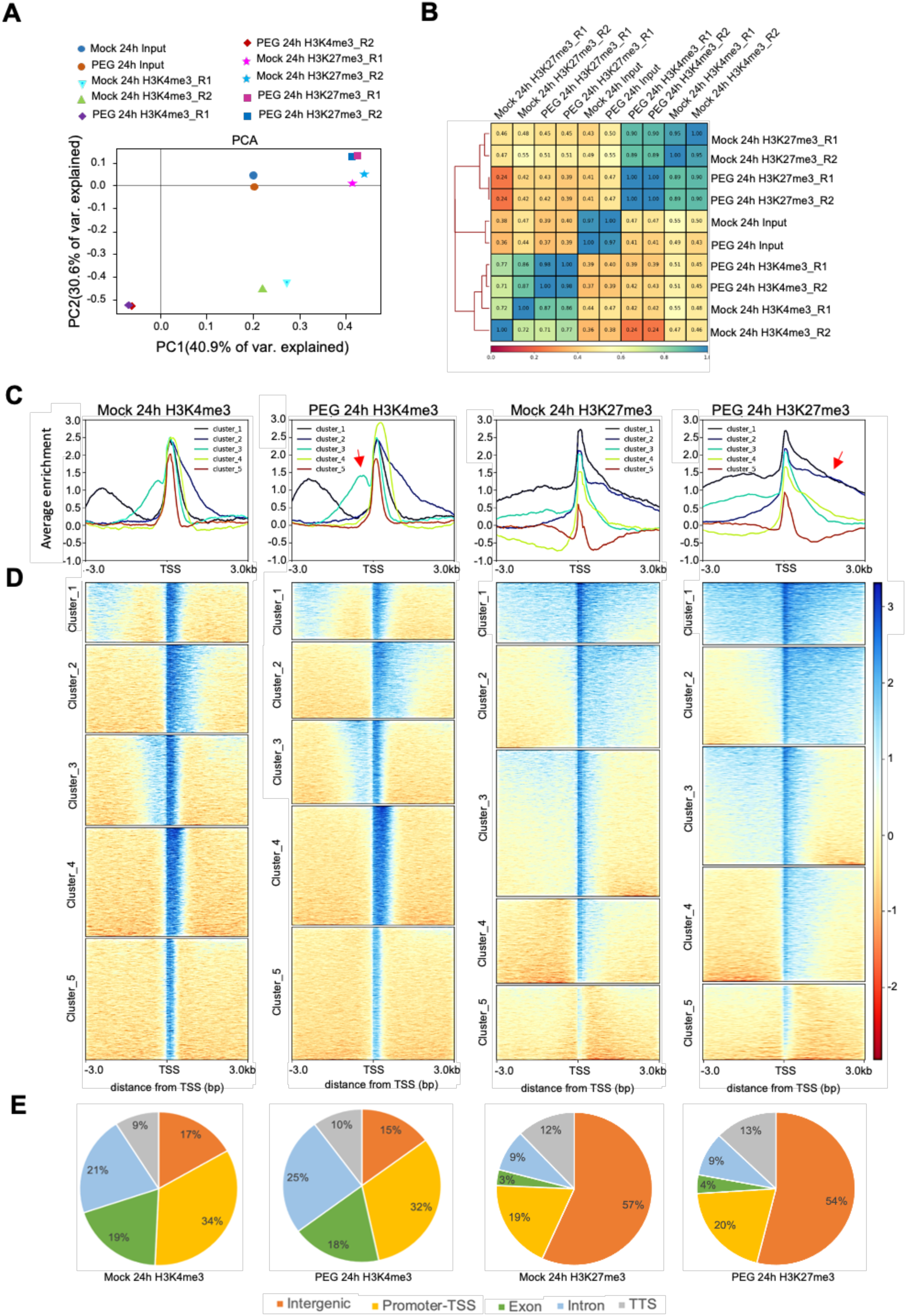
ChIP-Seq Analysis of Rapeseed in Response to PEG treatment. **A)** Principal Component Analysis (PCA) of Mock and PEG-treated datasets, illustrating distinct chromatin binding signatures for each condition. The percentage of variation is annotated on the PC1 and PC2 axes. **B)** Hierarchical clustered correlation matrix for Mock and PEG-treated replicates, based on ChIP-seq profiling data for H3K4me3 and H3K27me3 histone modifications. **C)** Average plot of ChIP-seq signals for H3K4me3 and H3K27me3 in Mock and PEG-treated samples. Meta-gene profiles were generated using the normalized sequencing density of H3K4me3 and H3K27me3. The X-axis shows the gene body as a percentage to standardize genes of different lengths, including regions 3 kb upstream and downstream of the gene. The Y-axis represents average values of H3K4me3 and H3K27me3. Stress-dependent change in the distribution of histonee marks across the regions are indicated by arrows. **D)** Heatmap of enriched ChIP-Seq signals for H3K4me3 and H3K27me3 in Mock and PEG-treated plants at 24 hours in five clusters. **E)** Genome-wide distribution of H3K4me3 and H3K27me3 displayed different genomic regions under PEG treatment and control conditions.

Subsequently we conducted a comprehensive assessment of genome coverage and ChIP-seq enrichment concurrently, employing fingerprint plot analysis. In the context of ChIP enrichment, the rightward deflection of the trace signifies the degree of enrichment observed. In this regard, all datasets exhibited ChIP enrichment when compared to the input, as shown in the fingerprint plot analysis (**Figure S12A, B**).

Our plot profile analysis revealed distinctive patterns in the distribution of H3K4me3 and H3K27me3 marks on rapeseed genome. Using a k-means clustering assay with H3K4me3 and H3K27me3-associated genes, we identified five clusters with distinct ChIP-seq read distributions for these histone modifications (**Figure 4C, D**). Clusters 1 and 3 exhibited broad H3K4me3 enrichment in both promoter and transcription start site (TSS) regions, while Cluster 2 showed H3K4me3 distribution in the gene body as well as in the TSS. Clusters 4 and 5 displayed intense H3K4me3 enrichment at the TSS site only. For H3K27me3 distribution, Cluster 1 genes had marks in the promoter, TSS, and gene body regions, whereas Clusters 2 and 4 showed H3K27me3 distribution in the gene body. Cluster 3 exhibited H3K27me3 enrichment in the promoter and TSS regions, and Cluster 5 restricted H3K27me3 marks to the TSS site only. Additionally, upon comparing PEG-treated samples with control samples, we observed subtle alterations in the plot profile trends for H3K4me3 marks. A slight increase in H3K4me3 marks was observed for Cluster 3 just before the TSS in the PEG-treated samples. Moreover, PEG treatment induced a broader distribution and heightened enrichment of H3K27me3 marks compared to control samples (**Figure 4C)**.

To visualize the distribution of marks across rapeseed genomes, we divided each genome into five annotated subregions: promoter-TSS (encompassing promoter and TSS), exons, introns, downstream TTS, and distal intergenic regions. Analysis of H3K4me3 and H3K27me3 distribution revealed subtle differences between these regions under both analyzed conditions (**Figure 4E**). Compared to the control, PEG-treated samples showed a reduction of approximately 1% in H3K4me3 marks in intergenic and exon regions, and 2% in promoter-TSS regions. Conversely, there was an increase of 4% in intronic and 1% in TTS regions for H3K4me3 marks. For H3K27me3, intergenic regions exhibited a 3% decrease, while promoter and TTS regions showed a 1% increase in distribution in PEG-treated samples, indicating potential mark-specific functional divergence.

To investigate the genomic distribution patterns of H3K4me3 and H3K27me3 marks in chromatin of PEG-treated rapeseed, we employed differential binding (Diffbind) analysis on mapped reads. Our analysis revealed 36,998 enriched regions associated with H3K4me3 and 37,541 with H3K27me3. Applying stringent criteria (False Discovery Rate ≤ 0.05 and Log2fold change ≥ 1) to normalize control H3K4me3 enriched regions, we identified 1,603 regions with differential H3K4me3 binding (**Figure 5A, Figure S12C-D**) and 1,426 regions with differential H3K27me3 binding (**Figure 5A, Figure S12E-F**). Enriched H3K4me3 regions encompassed 1,157 genes, with 976 gaining and 181 losing binding, alongside 272 transposons (227 gaining and 45 losing binding), and 174 non-annotated regions (153 gaining and 21 losing binding) (**Figure 5B**). H3K27me3 marks were notably associated with transposon regions, with 250 gaining and 519 losing marks, while 330 genes displayed altered binding patterns (169 gaining and 161 losing marks), and 327 non-annotated regions showed H3K27me3 peaks (**Figure 5C**). Representative transcripts illustrated the distribution patterns of H3K4me3 and H3K27me3 marks on mRNA transcripts of *BnP5CSA.3* and *BnPOX* genes **(Figures 5D and E)** and transposable element (TE) transcripts **(Figures 5F and G)**. This analysis allows for a comparative evaluation of the impact of chromatin mark distribution on genic and transposon regions, as well as on the expression levels of transcripts originating from these regions, between control and PEG-treated samples.

**Figure 5:**
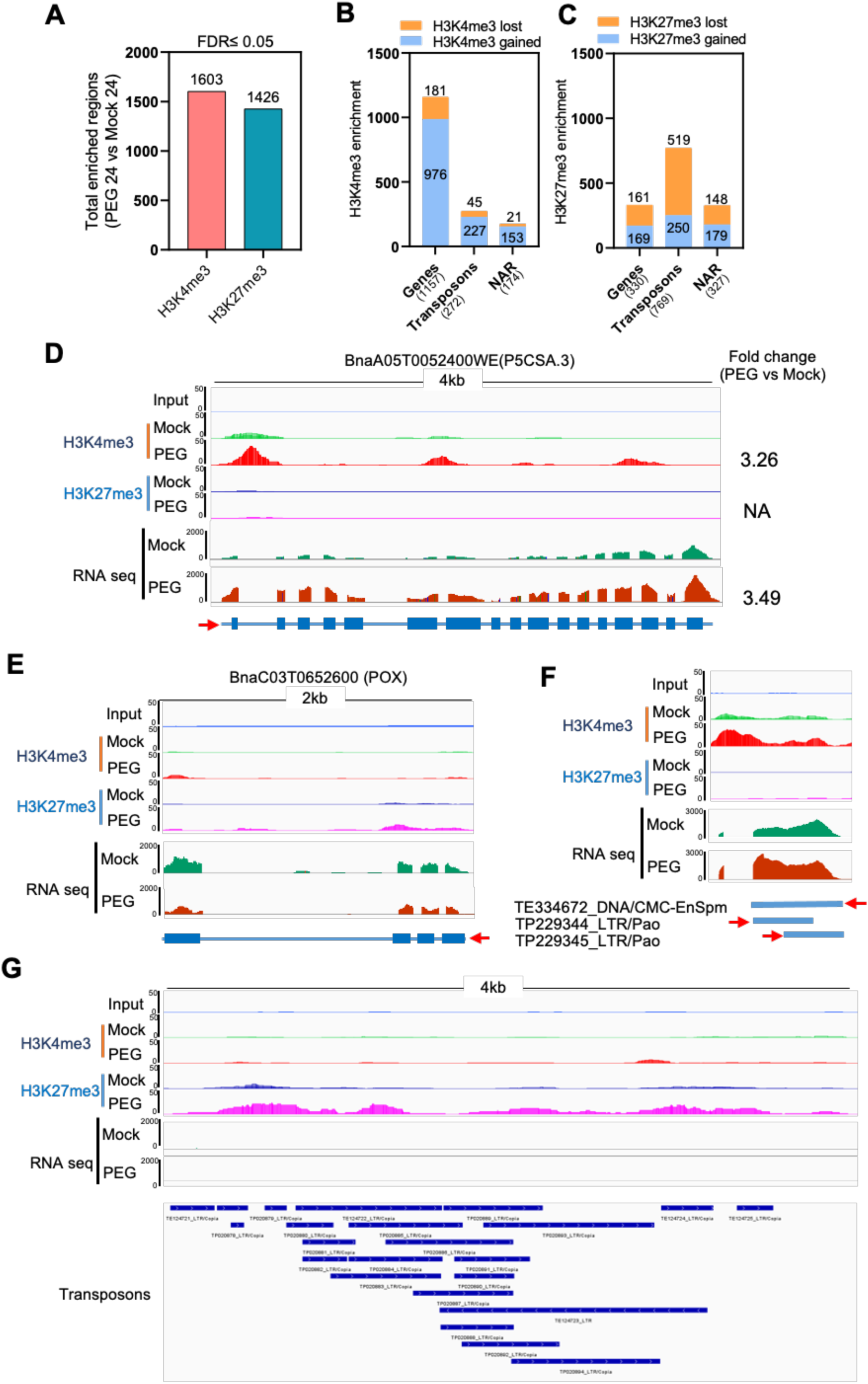
ChIP-Seq for Active Histone Marks H3K4me3 and H3K27me3. **A**) Bar graph illustrating the total enriched regions for H3K4me3 and H3K27me3 marks in PEG-treated samples compared to control (mock) samples. **B, C**) Stacked bar plots depicting the genomic distribution of H3K4me3 (**B**) and H3K27me3 (**C**) marks, showing identified gained and lost sites in rapeseed under PEG treatment, categorized by their association with genes, transposons, and nonannotated regions (NAR). **D, E, F**) ChIP-Seq and RNA-Seq tracks for the drought-responsive BnP5CSA.3 gene (**D**), BnaC03T0652600 (POX) gene (**E**), and a region encoding TEs (**F**). The Y-axis represents normalized read counts of H3K4me3, H3K27me3, and transcript reads (RPKM) within the locus (X-axis), with the direction of transcription indicated by arrowheads. **G**) ChIP-Seq tracks of genomic regions enriched with H3K27me3 that encode various TEs. The Y-axis represents normalized read counts of H3K27me3 within the locus (X-axis).

GO analysis of genes associated with H3K4me3, which exhibited gained binding, highlighted their predominant involvement in categories such as protein binding, nucleotide binding, heterocyclic compound binding, and processes related to methylation and macromolecule modification (**Figure S13A**). Conversely, genes showing loss of H3K4me3 marks were primarily associated with transferase activity and ion transport (**Figure S13B**). Further analysis revealed distinct trends for genes with gained H3K27me3 marks, which were enriched in hydrolase activity, response to stress, and defense responses. In contrast, genes losing H3K27me3 marks were associated with DNA binding, transcriptional activity, and metabolic processes (**Figure S14A-B**).

Our focus extended to understanding the distribution of H3K4me3 and H3K27me3 marks across transposable element (TE) regions under PEG stress. Enrichment of H3K4me3 and H3K27me3 was predominantly observed within retrotransposons, particularly those of the LTR-Copia, LTR-Gypsy, and LINE families, whereas such marks were less pronounced in DNA transposon regions (**Figure S15A and S16**). Certain regions of transposable elements (TEs) exhibit enrichment for both H3K4me3 and H3K27me3 marks, while a few regions are restricted to either H3K4me3 or H3K27me3 marks. Our results revealed that the level of enrichment is different for these marks **(Figure S15B)**.

### 5. Histone modifications associated with gene expression changes in rapeseed

To further investigate whether the observed gene expression patterns in the transcriptomic data correlate with histone mark distributions, we conducted a comparative analysis of RNAseq and ChIP-seq profiles. Among the 1,179 genes upregulated after 24 hours of PEG treatment, 58 genes exhibited an increase in the active H3K4me3 mark at the 5′-end of genes (downstream of the transcription start site). The IGV visualization of the three gene representations, BnaA09G0092200WE (*BnRAB18*), BnaC05G0053100WE (*BnPP2C*), and BnaC02G0218900WE (*BnH2A7*), showed upregulation upon PEG treatment and enrichment with H3K4me3 marks, as depicted in **Figure S17**. Conversely, among the 2,200 genes downregulated under the same conditions, 6 genes showed a gain and 9 genes showed a loss of H3K4me3 marks (**Figure 6A**). To better understand the relationship between increased H3K4me3 binding and enhanced gene expression, we performed scatter plot analysis for the 58 genes that showed increased H3K4me3 marks and were upregulated after 24 hours of PEG treatment. Plotting the Log2fold change in gene expression on the x-axis against the Log2fold increase in H3K4me3 binding on the y-axis revealed a noticeable correlation, suggesting that in these genes augmented H3K4me3 binding correlates with increased gene expression in response to PEG-induced stress (**Figure 6B**). Additionally, a heatmap illustrated the temporal and spatial expression patterns of these 58 genes with increased H3K4me3 marks (**Figure S18**).

**Figure 6:**
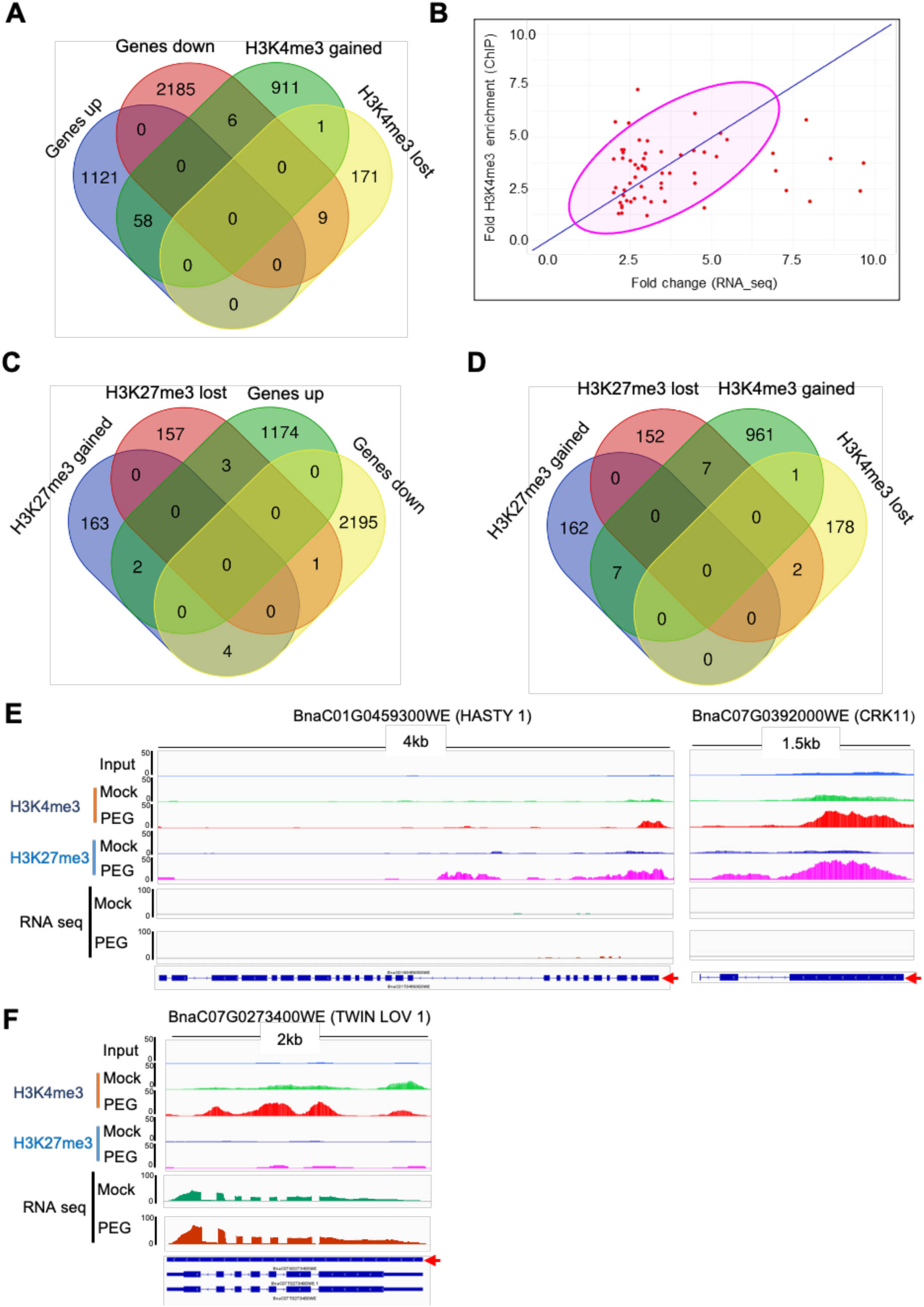
Integration of Rapeseed ChIP-Seq and RNA-Seq Data. **A)** Venn diagram illustrating the overlap between H3K4me3 targets and differentially expressed genes (DEGs) in response to 24 hours of PEG treatment. **B)** Scatter Plot visualization of the relationship between RNA-Seq and ChIP-Seq analysis for the 58 PEG-induced genes that gained H3K4me3 marks. Each point represents a gene, showing the correlation between changes in chromatin modification and gene expression. **C)** Venn diagram illustrating the overlap between H3K27me3 targets and DEGs in response to 24 hours of PEG treatment. **D)** Venn diagram displaying the intersection between genomic loci marked by H3K4me3 and H3K27me3, revealing genes that exhibit both active and repressive histone modifications. **E)** ChIP-Seq and RNA-seq tracks showing H3K4me3 and H3K27me3 bivalent chromatin marks at two representative genes (BnaC01G0459300WE: HASTY 1, BnaC07G0392000WE: CRK11) that gained both marks upon PEG treatment. **F)** ChIP-Seq and RNA-seq tracks of the gene TWIN LOV 1 (BnaC07G0273400WE) showing differential enrichment of H3K4me3 chromatin marks upon PEG treatment.

Many of the PEG-induced genes with increased H3K4me3 marks are implicated in stress responses, and includes transcription factors (ABF3/ABI5-like BnaA03G0547000WE, BnaC07G0458800WE, GBF3-type BnaA05G0007600WE, HSFA6A BnaA02G0281800WE), proteins involved in signal transduction (ABI2/PP2C protein phosphatase BnaA10G0055300WE, BnaC08G0480600WE, BnaC09G0377500WE, RLK-like kinase BnaC07G0252200WE), HSP proteins and chaperons (HSP40 BnaC07G0470800WE, BCS1- like mitochondrial chaperon BnaC09G0522300WE), E3 ubiquitin-protein ligase (BnaC08G0125900WE), and several dehydrin and LEA-type proteins (LTI65 BnaC02G0177500WE, RAB18 BnaA09G0092200WE, BnaC09G0109500WE, GMPM1/LEA-like BnaC09G0553200WE). Two PEG-induced P5CSA genes (one of them with spliced isoforms) could be identified among those which had enhanced H3K4me3 marks (BnaA05G0052400WE, BnaC04G0070800WE and BnaC04G0070800WE.1) suggesting that several genes controlling the rate limiting step in proline biosynthesis are under substantial epigenetic regulation. Expression profile and H3K4me3 enrichment of selected stress- responsive genes are shown **Figure S18**.

Regarding H3K27me3 marks, 2 genes exhibited increased and 3 genes showed decreased H3K27me3 marks among the upregulated genes. Conversely, among the 2,200 genes that were downregulated in response to PEG-induced stress, 4 genes gained and 1 gene lost the H3K27me3 mark (**Figure 6C**). Regulation and histone modifications, such as methylation, acetylation, phosphorylation, and SUMOylation, may also play a role in the regulation of gene expression in response to abiotic stresses like drought, high salinity, heat, and cold in plants (Liu et al., 2012; Fang et al., 2014; Kim et al., 2015; Wang et al., 2015). Our study indicates that, in addition to H3K4me3 and H3K27me3, other epigenetic marks may contribute to the regulation of gene expression in response to osmotic stress in rapeseed. Although H3K27me3 and H3K4me3 modifications are traditionally considered counterbalancing and mutually exclusive (Schwartz and Pirrotta, 2007; Bouyer et al., 2011; Roudier et al., 2011), our findings suggest otherwise for the response genes examined in this study. First, the levels of H3K27me3 and H3K4me3 were independent of each other’s presence. H3K27me3 levels remained constant regardless of the transcriptional status of most stress-responsive genes, whereas H3K4me3 levels showed a dynamic correlation with gene transcription, indicating that these two histone marks do not counterbalance each other. Second, the presence of high levels of H3K27me3 at a number of stress-responsive genes did not inhibit the accumulation of H3K4me3 during active transcription of these genes. Similar results were observed in Arabidopsis H3K4me3 and H3K27me3 marks function independently and are not mutually exclusive at the dehydration stress-responding memory genes (Liu et al., 2014a). A study on Arabidopsis in response to cold stress suggests that H3K4me3 and H3K27me3 differential methylation occur on distinct gene sets. Specifically, H3K4me3 differential methylation predominantly targets stress-responsive genes, while H3K27me3 differential methylation is primarily associated with developmental genes (Zeng et al., 2019).

Research on cold stress responses of Arabidopsis and potato unveiled a specific chromatin state known as bivalency, characterized by the coexistence of two histone marks: H3K4me3, which activates genes, and H3K27me3, which silences them (Faivre and Schubert, 2023). Further studies revealed that this chromatin state is not exclusive to cold-inducible genes; it also adorns numerous stress-responsive genes that can be reversibly silenced. This bivalency may prepare these genes for expression by keeping them in an accessible chromatin conformation facilitating transcriptional plasticity (Faivre and Schubert, 2023). To identify possible bivalent targets in rapeseed in response to PEG-induced stress, we analysed chromosomal regions enriched with H3K4me3 and H3K27me3. We could identify 7 genes that gained both of these histone marks, 7 genes gained H3K4me3 while losing H3K27me3 and 2 genes lost both histone marks following PEG treatment (**Figure 6D**). These results confirmed that co-occurrence of H3K4me3 and H3K27me3 bivalency happens in rapeseed in response to PEG-induced stress with low frequency.

To visualize the differential enrichment of H3K4me3 and H3K27me3 bivalent marks, we used IGV to examine two representative genes: BnaC01G0459300WE (*BnHASTY1-like*), BnaC07G0392000WE (*BnCRK11*), and BnaA05G0278200WE (**Figure 6E**). HASTY 1 is known for its role in miRNA biogenesis and is a candidate for mediating the export of plant miRNAs from the nucleus to the cytoplasm. Cysteine (C)-rich receptor-like kinases (CRKs), such as CRK11, are crucial for disease resistance and cell death in plants. The epigenetic regulation of these genes under stress conditions has not been previously explored. Our study revealed that both H3K4me3 and H3K27me3 bivalent marks were present on these genes. However, there was no significant increase in their expression levels under PEG stress compared to control conditions. The functional significance of these bivalent marks on these gene regions remains unclear. Additionally, we observed that the genomic region coding for BnaC07G0273400WE (TWIN LOV 1) exhibited both gain and loss of H3K4me3 marks in different regions following PEG treatment. Notably, transcript levels of TWIN LOV 1 increased under stress conditions compared to the control (**Figure 6F**). The role of LOV1 domain-containing proteins in plant stress responses is still not well understood.

In our study, we compared the transcriptomic and epigenetic profiles of B. napus with those of its diploid progenitors. In the *B. napus* Westar variety, 100,194 transcripts were identified, with 43,246 (43.16%) derived from the A-subgenome, 50,845 (50.75%) from the C-subgenome, and approximately 6,103 (6.09%) originating from scaffold sequences (**Figure S19A**). The expression levels of 24,508 (about 56%), 27,969 (about 55%), and 2,353 (about 39%) genes were actively expressed in the A, C, and scaffold subgenomes, respectively (**Figure S19B**). Our transcriptomic analysis identified 5,473 and 3,379 differentially expressed genes (DEGs) in *B. napus* under PEG stress at 6 hours and 24 hours, respectively (**Figure 3C-D**). The results revealed that transcripts from both the A and C-subgenomes exhibited a similar distribution of expression levels at both 6 and 24 hours post-PEG treatment (**Figure S19C-D**). Notably, regions derived from the C-subgenome showed a higher enrichment of H3K4me3 marks compared to those from the A-subgenome (**Figure S19E**). Additionally, a greater number of regions with H3K27me3 loss were detected in the C-subgenome (**Figure S19F**).

### Epigenetic regulation of rapeseed P5CS genes

Rapeseed genome is more complex than Arabidopsis, which is reflected by the higher numbers of genes which encode key enzymes in proline metabolism. Comprehensive analysis of transcriptome and ChIP data suggests that several of these rapeseed genes are regulated by osmotic stress. RNA-seq analysis indicates that most *BnP5CSA* genes and two out of the four *BnP5CSB* genes are induced by PEG treatment, with great variation in their transcript abundane (**Figure 7D, Figure S20**). Transcriptional activation correlated with the enrichment of the H3K4me3 activation mark in *BnP5CSA.3, BnP4CSA.5, BnP5CSB.4* genes (**Figure S20, Table S6**). Three out of the identified 14 P5CR genes had detectable transcripts, which was not significantly infuenced by PEG. Fast activation of *BnPDH1.1* and *BnPDH1.4* was found after 6 hours of PEG, while expression of *BnPDH2.1* was enhanced only after 24 hours of PEG (**Table S6**). Two out of the five *BnP5CDH* genes were expressed and could be slightly induced by PEG (*BnP5CDH.2, BnP5CDH.5*) (**Table S6**). None of these genes had differences in H3K4me3 or H3K27me3 marks, suggesting that only the stress-induced *BnP5CS* genes are under such epigenetic control (**Table S6**).

**Figure 7:**
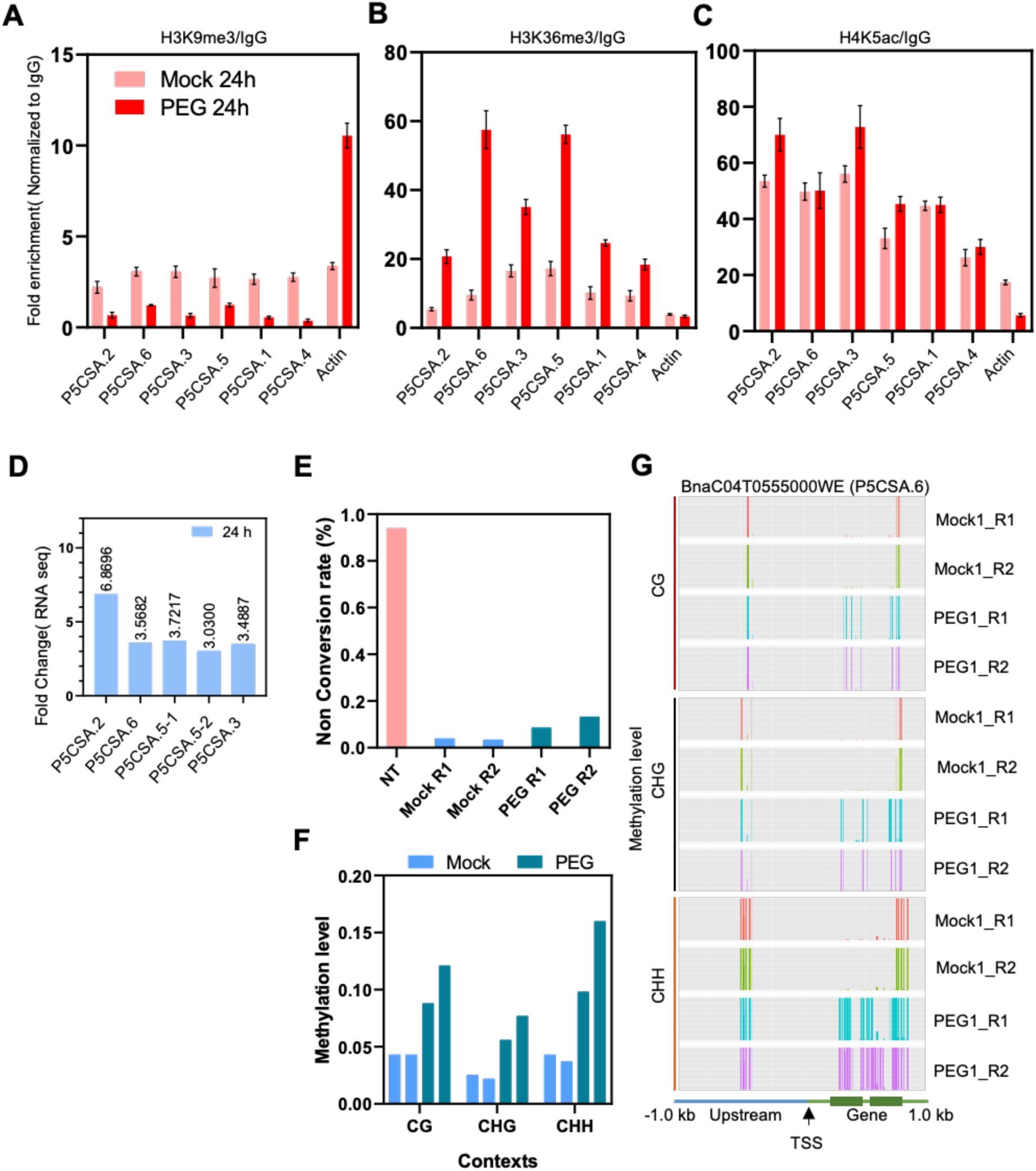
Epigenetic Regulation of P5CSA in Response to PEG Treatment. **A, B, C)** Differential enrichment patterns for H3K9me3 **(A),** H3K36me3 **(B),** and H4K5ac **(C)** at rapeseed *BnP5CSA* genes in response to 24 hours of PEG treatment, obtained with ChIP-qPCR analysis. **D)** Enhancement of rapeseed *BnP5CSA* expression in response to 24 hours of PEG treatment. **E)** Non-conversion rate of rapeseed DNA isolated from 24-hour Mock and PEG-treated samples. **F)** Average DNA methylation levels at *BnP5CSA* loci in different cytosine contexts (CG, CHG, and CHH) in response to 24 hours of 20% PEG treatment. **G)** DNA methylation profile at the P5CSA.6 region (BnaC04T0555000WE), including 1 kb upstream and 1 kb downstream from the transcription start site (TSS), in 24-hour Mock and PEG-treated samples. High-resolution map of DNA methylation patterns obtained from targeted bisulfite sequencing.

To extend our studies to other histone modifications in these genes, activation marks such as H3K36me3 and H4K5Ac, as well as the repression mark H3K9me3 was tested across BnP5CSA genes. ChIP was performed using anti-H3K36me3, anti-H4K5Ac, and anti- H3K9me3 antibodies, followed by qPCR analyses with primers targeting the TSS regions (**Table S2**). IgG ChIP and Actin genes were used as controls. The ChIP-qPCR data revealed a reduction in H3K9me3 across all BnP5CS genes upon PEG treatment compared to control (**Figure 7A**). The level of the H3K36me3 activation mark was significantly higher in the PEG-treated samples compared to controls (**Figure 7B**), suggesting that this chromatin modification may promote the expression of BnP5CSA genes. Among the rapeseed P5CS genes, BnP5CSA.1, BnP5CSA.3, and BnP5CSA.5 showed enrichment of the H4K5Ac mark upon stress, while the other P5CS genes did not exhibit significant differences between control and PEG treatment (**Figure 7C**). Notably, these trends were not observed in the Actin gene. Histone acetylation generally promotes gene expression. A study on rice seedlings subjected to drought treatment showed a considerable increase in the acetylation levels of total histone H3, as well as specific sites including H3K9, H3K18, H3K27, and H4K5 (Fang et al., 2014). In addition, our transcriptome analysis revealed differential expression of various histone acetyltransferases (HATs) and histone deacetylases (HDACs) under stress conditions, which can modulate the balance of histone acetylation (**Figure S6**).

### 6. PEG-Induced Hypermethylation in a Rapeseed P5CSA Gene

DNA methylation is a crucial epigenetic modification that plays a pivotal role in regulating numerous genes involved in stress responses. We were particularly interested in the potential influence of DNA methylation on the expression of stress-induced P5CS genes. To investigate this, we selected the BnaC04G0555000WE gene, which encodes P5CSA.6 and shows significantly elevated expression in PEG-treated plants at the 24-hour timepoint (**Figure 7D, Table S6**). For targeted bisulfite sequencing, we designed specific primers for this gene, covering 1 kilobase upstream and 1 kilobase downstream of the transcription start site (**Table S5**). Targeted bisulfite sequencing was then performed on both PEG-treated and control samples. The non-conversion rates of the samples are depicted in **Figure 7E**, with PEG-treated samples showing a slightly higher non-conversion rate compared to control samples. Our methylation analysis using Bismark tool revealed a PEG-dependent increase in methylation levels across all sequence contexts, including CG, CHG, and CHH, within the gene body (**Figure 7F**). Hypermethylation was particularly notable within the gene body regions. Although some methylation was detected at -0.5 kb in the upstream region, no PEG-dependent change was observed in the promoter of this gene (**Figure 7G**). These data suggest that enhanced DNA methylation in the transcribed region of the P5CSA.2 (BnaC04G0555000WE) gene may contribute to increased expression in osmotically stressed rapeseed plants.

## Discussion

### Physiological and molecular consequences of PEG treatment in rapeseed

Our study provides important insights into the complex responses of rapeseed to drought stress, revealing significant changes in gene expression and epigenetic regulation on genomic scale. The dynamic nature of these adjustments is essential for plant survival under fluctuating environmental conditions, with drought being a critical abiotic stressor that adversely affects plant growth and crop yields. Previous studies, predominantly made with model plants, have emphasized the role of epigenetic mechanisms, such as DNA methylation and histone modifications, in plant responses to abiotic stresses (Thiebaut et al., 2019, Nunez-Vazquez et al., 2022, Sun et al. 2022). However, information on the specific roles of chromatin modifications in non-model plants in water-restricted conditions is scarce. Transcriptional responses of salt and drought-treated rapeseed has been investigated by Wang et al. (2019), although the experimental conditions in that study were quite different. Through the integration of ChIP-seq and RNA-seq, we have identified gene sets in rapeseed that respond to osmotic stress, highlighting the epigenetic regulation involved.

PEG-induced osmotic stress can mimic water-restricted environment which generates physiological and developmental responses similar to water shortages during drought. PEG treatment can be controlled more precisely than water withdrawal, and has therefore been employed to characterize tolerance to drought (Verslues et al., 2006). In rapeseed, we observed substantial morphological, physiological, and biochemical changes in response to PEG, indicating the efficiency of the treatment. One notable response was the accumulation of proline, a common physiological response to osmotic and salinity stress. Proline accumulation is a sensitive indicator of stress condition, which often (but not always) correlates with tolerance to drought or salinity. Proline functions as an osmolyte, ROS scavenger, and molecular chaperone, playing a crucial role in various cellular processes, including osmoregulation, redox balance, and the maintenance of cellular homeostasis, and influencing gene expression(Szabados and Savouré 2010; Patriarca et al. 2021, Alvarez et al., 2022). In rapeseed leaves enhanced proline levels were observed after 24 h of PEG treatment, followed by accumulation up to 7 days of stress (**Figure 1**). In a transcriptomic study, Wang et al. (2019) reported around two-fold proline accumulation in whole rapeseed seedings only after three hours of PEG treatment. Our results contradicts to those results, as no difference in proline levels could be observed in rapeseed leaves even after 6 hours or in stems up to 48 hours of PEG treatment (proline accumulation in roots was negligible, **Figure 1, S1**).

Enhanced ROS levels and lipid peroxidation rates indicated the damaging effect of PEG on cellular structures (**Figure 1**). Photosynthesis was also affected by PEG treatment. We observed changes in several photosynthetic parameters under stress, including ECSt, Fv/Fm, NPQt and PhiNPQ as well as increased leaf temperature which could be the consequence of excess energy produced by NPQ and stomata closure which prevented evaporation (**Figure 1**). Drop in Fv/Fm ratio indicated reduction of PSII’s maximum photochemical efficiency, while enhanced ECSt is likely due to a heightened transthylakoid proton gradient and reduced ATP synthesis and carbon assimilation of PEG-treated plants (Buchert et al., 2021; Hikosaka and Tsujimoto, 2021). Profound changes in all these physiological parameters confirmed the detrimental effect of PEG treatment on rapeseed plants, suggesting that the system is suitable for genome-wide molecular studies.

Molecular responses to PEG-induced osmotic stress was monitored by testing the expression of selected genes including *BnP5CSA, BnP5CSB, BnRAB18, BnRD29A, BnRD29B, BnNCED3, BnRD22,* and *BnDREB2A*, whose Arabidopsis homologs are known to respond to osmotic stress (Strizhov et al., 1997; Sakuma et al., 2006; Matsui et al., 2008; Takahashi et al., 2018). Transcript levels of these genes were elevated in PEG-treated rapeseed plants (**Figure 2**), indicating that the osmotic stress applied was efficient to stimulate the signaling pathways, which were known to control the activities of these target genes (Matsui et al., 2008; Todaka et al., 2015; Takahashi et al., 2018). The drought-induced *NCED3* controls the rate-limiting step of ABA synthesis as described in Arabidopsis (Iuchi et al., 2001; Finkelstein, 2013). Upregulation of *BnNCED3* in PEG-treated rapeseed indicated enhanced ABA biosynthesis and ABA content, which could be responsible for the observed high transcript levels of ABA-responsive genes such as *BnP5CSA, BnRAB18, BnRD29A* and *BnRD29B* (**Figure 2**).

### RNAseq analysis reveals large-scale changes in transcript profiles of PEG-treated rapeseed

In order to identify gene sets that respond to osmotic stress in rapeseed leaves, transcript profiling was made after short (6 hours) and extended (24 hours) periods of PEG treatment. Number of differentially expressed genes (DEGs) in our experiment (**Figure 3, Figure S2**) was comparable to earier studies, which were obtained on drought-stressed rapeseed plants (Liu et al., 2015; Wang et al., 2017) or after salt treatment (Yong et al., 2014). Lower number of differentially expressed rapeseed genes were identified by Wang et al. (2017), who defined 169 DEGs to be crucial in response to drought. Our data however indicates that much higher number of genes are modulated by high osmotic conditions, as we found 5473 DEGs after 6 hours and 3379 DEGs after 24 hours of PEG treatment (**Figure 3**). An earlier study reported 913 DEGs in rapeseed after short time (3 hours) of PEG treatment (Wang et al. 2019). In that study ROS accumulation and oxidative damage was not observed and proline accumulation was meager, suggesting that the treated plants experienced minimal stress and their physiological status was not altered significantly. In our experimental conditions the osmotic stress was more robust, as reflected by the significantly altered physiological parameters and the high number of DEGs observed in RNAseq experiment.

Gene Ontology (GO) classification showed that a number of stress-responsive genes, signal transduction pathways, metabolic processes, and photosynthetic activities are altered in rapeseed upon osmotic stress. Specifically, genes responsible for DNA binding transcription factor activity, chromatin binding, transcription regulation, and stress response were upregulated, while genes involved in photosynthetic activity and metabolic processes were downregulated under osmotic stress. Importantly, genes such as *BnP5CS, BnRAB18, BnNCED3, BnRD22,* and *BnPOX* exhibited higher transcript levels, while most *BnProDH* and *BnSOD* genes showed lower transcripts upon osmotic stress. Transcription factors (TFs) are crucial regulatory proteins that bind to specific DNA sequences, thereby controlling the transcription of target genes. In plants, various classes of TFs are activated in response to drought stress, orchestrating complex gene expression changes to confer stress tolerance. Our study observed varying trends for different transcription factor families, with drought-specific transcription factors such as DREB, ABI/ABF, bZIP, HD-ZIP, MYB, NAC, and MYC showing elevated transcription levels (**Figure S9**). Identification of such transcription factors can facilitate the establishment of gene regulatory networks controlling responses to osmotic stress and drought.

Besides transcription factors, expression of a number of genes encoding chromatin regulators were modified by PEG treatment (**Figure S10**), suggesting that chromatin structure of rapeseed might be different in such conditions. Altered chromatin structure, can influence accessibility of DNA to transcription factors and RNA polymerases in stress conditions (Luo et al., 2012; Bhadouriya et al., 2021). Transcripts of various such genes were elevated in PEG-treated plants including *BnDRM2, BnSUVH6, BnHDAC, BnJMJ, BnRDR5, BnATX1,* and *BnSUVH1* factors, while numerous others had reduced transcript abundancy such as *BnMET1, BnCLASSY, BnHAC1, BnROS1,* and *BnDME* (**Figure S10**). Altered abundance of such chromatin regulators could modify the delicate balance between methylated and unmethylated DNA, and methylated or acetylated histones, which is crucial to control adaptive responses in plants (Penterman et al., 2007; Torres and Fujimori, 2015).

### Osmotic stress generates large-scale modifications of epigenetic marks in rapeseed

Numerous reports describe extensive DNA and histone modifications that modulate gene expression in plants under stress conditions. Such modifications can be reset to the basal level after recovery or sustained to generate stress memory which can even be transmitted to progenies (Chinnusamy and Zhu, 2009; Lämke and Bäurle, 2017). To reveal the consequences of PEG-induced osmotic stress on chromatin structure, a genome-wide analysis of histone modifications was performed. The role of histone methylation in regulating phenotypic responses to environmental changes in model and cultivated species has received increased attention in recent years (Shi et al., 2023; Abdulraheem et al., 2024). However, the regulatory landscape and functional relevance of histone modification in *Brassica napus* against drought stress remain poorly understood. We used a quantitative, single-base-resolution technique (ChIP-seq) to profile genome-wide patterns of H3K4me3 and H3K27me3, two key histone modifications involved in stress-induced gene activation and repression, respectively (Kim et al., 2015; Shi et al., 2023). H3K4me3 mainly located within genic regions, whereas H3K27me3 significantly enriched within intergenic regions including TE elements (**Figure 4**). Our analysis revealed that in the genes of PEG-treated plants nearly six times more H3K4me3 marks could be identified when compared to H3K27me3 marks (976 versus 169), while the amount of lost H3K4me3 and H3K27me3 marks were similar in genes (181 versus 161) (**Figure 5**). Such difference suggests that influence of H3K4me3 on gene activation is dominant over the effect of H3K27me3 in osmotically stressed rapeseed plants. Similar trend was reported in Arabidopsis, where H3K4me3 correlated with drought-dependent activation of a set of stress-induced genes (Kim et al., 2008). H3K4me3 and H3K27me3 often act antagonistically in plants exposed to environmental stress, with H3K4me3 acting as an active mark and H3K27me3 as a repressive mark (Kim et al. 2015). GO analysis revealed that genes associated with transcription regulation, methylation, and metabolic processes were enriched with H3K4me3 marks while genes responsible for cell cycle processes and ion transport lost H3K4me3 marks upon PEG treatment (**Figure S13**).

Differential accumulation of the H3K27me3 mark was observed in a small number of genes in rapeseed under PEG stress. Genes encoding TRANSPARENT TESTA-12 like, translation initiation factor IF-3-like, AP2-like ethylene-responsive transcription factor, autophagy-related protein, MADS-box protein CMB1-like, and cytosolic sulfotransferase 16-like gained the H3K27me3 mark, while genes like disease resistance protein TAO1-like, ethylene-responsive transcription factor 1A-like, and PIF3 lost the H3K27me3 marks upon PEG treatment (**Supplementary dataset 10**).

Despite generally distinct genomic locations and functions between the two histone modifications, we identified seven overlapping regions, possibly due to promoter bivalency (H3K4me3-H3K27me3) (Faivre and Schubert, 2023). Genes encoding for HASTY 1, WRKY transcription factor 19 (WRKY19), Cysteine-rich receptor-like protein kinase 11 (CRK11), Putative protease Do-like 14 (DEGP14) /PARK13, an F-box protein and several uncharacterized genes showed lower transcript abundance under PEG stress. Additionally, the gene encoding TWIN LOV 1 (BnaC07G0273400WE), a photoreceptor protein for blue light exhibited a different pattern of H3K4me3 distribution within the gene body (**Figure 6**). In the control condition, H3K4me3 was enriched at the TSS site of these genes, but under stress we observed redistribution of these marks in the gene body, which may have influenced transcript accumulation.

These results provide new insights into the chromatin-mediated epigenetic regulation of transcription in response to drought stress and highlight the need for further investigation into the functional implications of these epigenetic changes. Our ChIP analysis for H3K27me3 showed that this mark is mostly distributed in the TE regions, encoding for LTR/Copia, LTR/Gypsy, DNA/CMC-EnSpm, and LINE/L1 transposons (**Figure S16**). Further analysis of TE transcripts encoded by the H3K27me3 binding regions is needed to understand the influence of PEG stress on the function of this epigenetic mark on regulation of TE elements.

A detailed analysis of genes with significant changes in epigenetic marks revealed that 58 genes had significantly higher expression and enrichment with active H3K4me3 upon PEG stress, suggesting positive impact of the H3K4me3 mark on expression of these genes (**Figure S18**). Characterization of this gene set showed that many stress-related genes like *BnP5CS*, *BnRAB18, BnABI5,* Dehydrins, *BnRD23D,* Histone H2A7, RMA3-like, protein phosphatase 2C, and Heat stress transcription factor 6 (*BnHSF6*) had higher expression and were enriched with H3K4me3. Enriched H3K4me3 mark could contribute to enhanced expression of stress-responsive genes in PEG-treated rapeseed. Such epigenetic change can also facilitate adaptation to drought in field conditions.

The allotetraploid *Brassica napus* (AACC) originated from the hybridization of two diploid species, *Brassica rapa* (2n = 20, AA) and *Brassica oleracea* (2n = 18, CC). Previous studies have documented the phenomenon of asymmetric genomic expression in polyploids (Liu et al. 2014b). In *B. napus*, as an allopolyploid, the two subgenomes have undergone biased segregation following polyploidization, leading to asymmetric genome evolution (Bottani et al., 2018). This evolution is driven by a variety of complex factors, including the asymmetric loss of subgenomes, differential amplification of transposable elements and tandem repeats, and variations in DNA sequence and expression (Liu et al. 2014b). Such biased segregation has resulted in varying degrees of gene loss (Wendel et al., 2018), with the processes of polyploidization and gene loss altering the composition of gene families, potentially contributing to the observed morphological plasticity in Brassica species (Wang et al., 2011). In our study, we observed no significant differences in the distribution of DEGs between 6- hour and 24-hour PEG-treated samples, with both the A-genome and C-subgenome contributing equally to the response. However, epigenetic analysis of H3K4me3 and H3K27me3 revealed a greater number of regions in the C-subgenome that were altered under stress conditions. These findings suggest that the C-subgenome in *B. napus* may experience a higher frequency of epigenetic modifications and maintain greater nucleotide diversity compared to the A-subgenome in the Westar variety (**Figure S19**).

### Epigenetic control of P5CS genes in rapeseed during osmotic stress

P5CS is a rate-limiting enzyme for proline synthesis, and is encoded by two genes in Arabidopsis, *P5CS1* and *P5CS2* (Strizhov et al., 1997). *P5CS1* responds to drought and salt stress and is regulated by ABA-dependent and ABA-insensitive signalling pathways, while *P5CS2* has a rather constitutive expression (Yoshiba et al. 1995; Savouré et al. 1997; Székely et al. 2008). Mutations disrupting the *AtP5CS1* gene display salt hypersensitivity, confirming its importance in stress tolerance (Szekely et al., 2008). We have identified ten P5CS genes in rapeseed, grouped into two subfamilies, *BnP5CSA* and *BnP5CSB* (**Figure S20, Table S6**). Transcript analysis of individual P5CS genes was not possible with qRT-PCR technique due to high sequence identity. We could however differentiate between the *BnP5CSA* and *BnP5CSB* subfamilies by qRT-PCR, and showed that these genes were upregulated by PEG-generated osmotic stress (**Figure 2**). RNAseq analysis could give more precise information about the regulation of individual P5CS genes. Four out of the six *BnP5CSA* genes and two of the four *BnP5CSB* genes were significantly induced by PEG at both 6 h and 24 h treatments. (**Figure S20, Table S6**). The other P5CS genes were not or only moderately activated during PEG treatment. This scenario resembles to other plants such as Arabidopsis, which posses stress-induced and constitutive P5CS genes, the later with housekeeping function (Székely et al. 2008).

Comprehensive analysis of transcriptomic and histone modifications via ChIP shows a strong correlation between osmotic stress-induced gene expression and enrichment of epigenetic marks in several BnP5CS genes. ChIP-seq analysis revealed that regions encoding *BnP5CSA.3, BnP5CSA.5* and *BnP5CSA.4* genes exhibited significant enrichment of H3K4me3 upon PEG treatment (**Table S6**). Our results suggest that other epigenetic marks such as H3K9me3, H3K36me3, and H4K5Ac can also influence the activity of rapeseed P5CSA genes (**Figure 7**). H3K9me3 is a repressive mark known to suppress the expression of various stress-responsive genes (Lu et al., 2022). Our study revealed a reduction in H3K9me3 across all *BnP5CSA* genes under stress. Additionally, H3K36me3 levels were enriched at the transcription start site (TSS) of the *BnP5CSA* encoding regions, while H4K5Ac showed minimal influence on most *BnP5CSA* genes, except for slight differential enrichment in *BnP5CSA.2* and *BnP5CSA.3*. Several *BnP5CR, BnPDH1, BnPDH2* and *BnP5CDH* genes had detectable transcript level, some of them displayed PEG-dependent activation. None of these genes however had altered histone modifications (**Table S6**), suggesting chromatin methylation has no influence on their expression.

Transcript mapping of *BnP5CSA.5* in the RNAseq experiment could identify three transcripts of this gene (**Figure S20, Table S6**), suggesting that alternative splicing might regulate at least one *BnP5CS* gene in rapeseed. Intron-mediated alternative splicing was shown to regulate *P5CS1* activity in Arabidopsis (Kesari et al. 2012). Alternative splicing can therefore control the activity of P5CS genes in various Brassicaceae species. Further studies are needed to characterize the consequences and function of *BnP5CSA.5* alternative splicing in rapeseed.

DNA methylation is a hallmark epigenetic modification that plays a crucial role in regulating various biological processes, including genome stability, chromatin structure, gene expression, and silencing of transposable elements in extreme environments. In plant genes DNA methylation has different functions, depending on the position: in promoters it usually inhibits gene expression, while in exons CG methylation may enhance transription in a dose-dependent manner(Lei et al., 2015; Thiebaut et al., 2019; He et al., 2022; Sun et al., 2022). Our analysis focused on the gene body region of the PEG-induced *BnP5CSA.6* gene, which had enrichment in H3K36me3 marks upon osmotic stress (**Figure 7, Table S6**). We found hypermethylation across all contexts (CG, CHG, and CHH) within Exon 1 and Exon 2 in PEG-treated samples, indicating that gene body hypermethylation can influence expression of this gene. Recent reports suggest, that Gene body methylation (GbM) does not correlate with transcriptional repression, but rather is associated with elevated gene expression and enhanced plasticity of gene activation (Muyle et al., 2022; Williams et al., 2023). GbM, particularly hypermethylation, has been shown to influence gene expression under various stress conditions, implicated in stabilization and upregulation of transcription (Chinnusamy and Zhu, 2009; Muyle et al., 2022; Sun et al., 2022). GbM of *BnP5CSA.6* may contribute to its activation in PEG-treated rapeseed plants. The downregulation of *BnDME* and *BnROS1* DNA demethylases observed in the transcriptome data (**Figure S6**) might be the reason for the hypermethylation levels in the *BnP5CSA.6* gene under PEG-induced osmotic stress (**Figure 7**). In Arabidopsis increased H3K4me3 levels of the stress-induced *P5CS1* gene were observed in salt-treated plants and found to be maintained after stress recovery which may contribute to transcriptional memory and increased *P5CS1* expression in repeated stresses (Feng et al. 2016). These observation aligns with results obtained on other plant species. Drought stress in tomato roots resulted in decreased CHH methylation in regulatory regions but increased CHG and CHH methylation in coding regions of the drought-responsive gene ABSCISIC ACID STRESS RIPENING 2 (*SlAsr2*) (González et al., 2013). DNA hypermethylation in salt tolerant genotypes of rice was also observed, while sensitive genotypes exhibited demethylation (Feng et al., 2012). Such findings underscore the potential regulatory roles of gene body hypermethylation suggesting that it may contribute to stress-dependent alterations of gene expression (Muyle et al., 2022; Sun et al., 2022). GbM can be transmitted to progenies suggesting that it can play certain role in stress adaptation and (Muyle et al., 2022; Sun et al. 2022, Williams et al., 2023). Further research is however needed to unravel the precise mechanisms by which GbM influences gene expression in rapeseed and to determine its broader implications in stress tolerance and adaptation. Collectively, these results indicate that stress-induced proline accumulation in rapeseed is under complex epigenetic control which primarily regulates key P5CS genes controlling first and rate-limiting step of proline biosynthesis (Szabados and Savouré, 2010).

In conclusion, our study highlights the critical role of epigenetic modifications, particularly histone marks and DNA methylation, in mediating gene expression changes in rapeseed under PEG-induced osmotic stress. The generated expression and epigenetic profiles of rapeseed can facilitate the understanding of gene regulatory mechanisms on genomic scale in such allotetraploid crop with complex genome. These findings enhance our understanding of the molecular mechanisms underlying plant responses to abiotic stress and provide a foundation for developing strategies to improve crop resilience to adverse environmental conditions. Characterized epigenetic marks can be targets of genome editing with DNA modifiers or chromatin remodelers to edit epigenetic marks in crops like rapeseed and engineer gene expression profiles to improve drought tolerance (Selma and Orzáez2021; Seem et al., 2024).

## Materials and methods

### Plant material and growth conditions

Plants of *Brassica napus* var. Westar (Klassen et al., 1987) were grown on hydrophonic system using Hoagland nutrient solution as described(Wang et al., 2019). Plants were maintained and all treatments were performed in a growth chamber (Fytoscope SW, PSI, Czech Republic), under precisely controlled conditions. Illumination was provided with LED panels with photon flux of 160 μmol of photons m-2s-1 and a short-day light cycle (8 h light at 22°C / 16 h darkness at 20°C). Four-weeks-old plants were subjected to osmotic stress by replacing the standard Hoagland nutrient solution with solution supplemented by 20% PEG8000. Plants were allowed to recover from osmotic stress by replacing the PEG-containing solution with standard Hoagland solution. Leaf samples of PEG-treated and control plants were collected at various time points, specifically at 6 hours, 24 hours, 48 hours, 3 days, 5 days, 7 days, 10 days, and 15 days, immediately frozen in liquid nitrogen and stored until processing (workflow is show in Figure S1A). Experiments were repeated three times (biological replicates).

### Determination of proline content

Proline content was determined as described with minor modifications(Kovács et al., 2019). 50 mg of leaf tissue was ground with liquid nitrogen, 1 ml of 1% sulfosalicylic acid was added and mixed by vortex. The mixture was centrifuged at 13000 rpm for 10 min at 4°C, and 200 μl supernatant was mixed with 400 μl of 1.25 % ninhydrin reagent (ninhydrin dissolved in 80 % acetic acid). The samples were incubated in the dry block heater bath at 95°C for 30 min, and immediately placed on ice for several minutes. Proline content was determined by measuring the absorbance of the reaction product at 520 nm using Thermo Scientific, Multiscan Go Microplate Spectrophotometer. To draw the standard curve of proline, concentrations of 0.5, 0.25, 0.125, 0.025, 0.03125 and 0 mM proline were used as reference. The content of proline was measured with five technical replicates.

### Determination of malondialdehyde

Lipid peroxidation rates were measured by the thiobarbituric acid-reactive substances (TBARS) assay as described (Heath and Packer, 1968). 100 mg leaf tissue was homogenized in 1 ml of 0.1% trichloroacetic acid (TCA) containing 0.4% butylhydroxytoluene, centrifuged at 13,000 rpm for 20 minutes. 1 ml of 20% TCA containing 0.5% thiobarbituric acid (TBA) was added to 250 μl supernatant, mixed and incubated at 96 °C for 30 min. Absorbance was measured at 532 nm using a Multiskan GO microplate reader (Thermo Fisher Scientific) and calculated by subtracting the nonspecific absorbance measured at 600 nm. Malondialdehyde (MDA) concentration was calculated using the extinction coefficient ε532 = 155 mM−1 cm−1. Three biological replicates were made.

### Determination of photosynthetic parameters

Chlorophyll fluorescence measurements on light-adapted leaves were carried out directly in the growth chamber using a portable MultispeQ v1 device (PhotosynQ) controlled by the PhotosynQ platform software (Kuhlgert et al., 2016), using a red AL (200 μmol photons m−2 s−1) and a 500-ms saturation pulse (3000 μmol photons m−2 s−1).

### RNA Extraction and RT-qPCR Analysis

Total RNA was isolated using a GeneJET RNA Purification Kit (Thermo scientific) following the manufacturer’s protocol. One μg of DNase-treated RNA was used for cDNA synthesis using the High-Capacity cDNA Reverse Transcription Kit (Applied Biosystems). qRT-PCR was performed on 3 μl of 20x diluted cDNA templates using 5x HOT FIREPol EvaGreen qPCR Mix Plus (Solis Biodyne) in a final volume of 10 μl, and Bio-Rad CFX96 Touch Deep Well Real-Time PCR System. Mean values of Actin and GAPDH Ct were used as internal reference. Normalized relative transcript levels were determined using the 2^-ΔΔCt^ method (Livak and Schmittgen, 2001). Experiments were repeated with three biological replicates. Oligonucleotides used in this study are listed in **Table S1**.

### RNA-seq analysis

Total RNA was isolated using a GeneJET RNA Purification Kit (Thermo scientific) following the manufacturer’s protocol. Approximately 3μg of total RNA from each sample was subjected to the RiboMinus Eukaryote Kit (Qiagen, Hilden, Germany) to remove ribosomal RNA prior to the construction of the RNA-seq libraries. RNA-seq libraries were prepared using an RNA-seq Library Prep Kit for Illumina (New England Biolabs). The libraries were sequenced on an Illumina HiSeq 4000 platform using a SE150-bp read module. The raw sequencing reads were filtered and trimmed using the default parameters of Fastp (version 0.23.2) (https://academic.oup.com/bioinformatics/article/34/17/i884/5093234). The filtered clean data were then pseudo-aligned against the transcript models for *Brassica napus* var. Westar downloaded from (http://cbi.hzau.edu.cn/cgi-bin/rape/download_ext) using Kallisto (version 0.50.1). Differentially expressed genes (DEGs) were identified using DESeq2 (version 1.42.1 (http://bioconductor.org/packages/release/bioc/html/DESeq.html) with the criteria of q value (adjusted p value, Benjamini-Hochberg method) <0.05 and |log_2_ Fold Change (FC)| ≥2)(Love et al., l., 2014). Functions of the DEGs were investigated with Gene Ontology (GO) and Kyoto Encyclopedia of Genes and Genomes (KEGG) pathway enrichment analysis using topGO (version 4.4) and ClusterProfiler (version 4.12.2)(Alexa and Rahnenfuhrer, 2010; Wu et al. 2021), respectively. Significant GO terms and KEGG pathways were identified with the criterion of q value <0.05. Sequence data were deposited in the National Center for Biotechnology Information (NCBI) SRA database (Accession number: PRJNA1144338)

### ChIP-seq and ChIP-qRT PCR

ChIP assays were performed as previously described by Song et al. (2016) with minor modifications. In brief, 2 to 3 g of 30-day old plant leaves (Mock and PEG treated) were crosslinked with 1% formaldehyde for 20 min under vacuum, quenched with freshly prepared 2M glycine and ground into fine powder in liquid nitrogen. Chromatin was isolated and sheared into 200 to 500 bp DNA fragments by sonication. The sonicated chromatin was immunoprecipitated with one of the following antibodies (5 μg): Anti-trimethyl-Histone H3 (Lys4) (Merck, 07-473), Anti-Histone H3K27me3 (Active motif, 39155), Anti-Histone H3 (di methyl K9) (Abcam, ab1220), Anti-Histone H3 (tri methyl K36) (Abcam ab9050), Anti-Histone H4 (acetyl K5) (Abcam, ab51997) or Anti-Myc tag antibody (used as IgG negative control, Abcam, ab9E11) and with 25 μl of Dynabeads Protein G (Invitrogen, 10003D) for 12 h at 4 °C with rotation. The precipitated chromatin DNA was then purified by phenol– chloroform-isoamyl alcohol extraction and recovered by ethanol precipitation. The ChIP DNA was prepared for sequencing or qPCR. Two biological replicates were used for ChIP-seq, and three biological replicates were used for ChIP–qPCR. ChIP-IgG was used for normalising the values. the primers used for ChIP qPCR are listed in **Table S2**. Actin (BnaC09T0320800WE) was used as a negative control.

### ChIP-seq data analysis

At least 10 ng of ChIP DNA was used for library preparation. ChIP sequencing libraries were constructed using NEBNext Ultra II DNA Library Prep Kit (New England Biolabs Inc, E7103) following the manufacturer’s instructions. Constructed libraries were sequenced using Illumina NovaSeq 6000 and paired-end reads were obtained at Geninus Bowtie2 software (https://www.kr-geninus.com) (Langmead and Salzberg, 2012) and were used to align the sequencing reads of ChIP-seq to the *Brassica_napus* reference genome (http://cbi.hzau.edu.cn/rape/download_ext/westar.genome.fa) using default parameters. The peak in different conditions and differentially changed peaks were called by MACS software (Zhang et al., 2008). The nomodel parameter was set, and the d-value parameter was set at 200. The resulting wiggle files, which represent counts of ChIP-Seq reads across the reference genome, were normalized for sequencing depth by dividing the read counts in each bin by the millions of mapped reads in each sample and were visualized in the IGV genome browser. The Diffbind was used to Compute differentially bound sites between PEG-treated and control conditions from multiple ChIP-seq experiments using affinity (quantitative) data. For the analysis of histone modification profiles between the 24-hour PEG-treated samples and the mock-treated samples, the Galaxy platform was employed (<u>usegalaxy.eu</u>). Sequence data were deposited in the National Center for Biotechnology Information (NCBI) SRA database (Accession number: PRJNA1144218).

### Targetted DNA methylation

Primers for the targeted DNA methylation were designed using Bisulphite-primer-seeker tool (https://zymoresearch.eu). DNA from the rapeseed was isolated using DNA extraction kit Nucleon PhytoPure (Merck). Bisulphite conversion of DNA was carried using the kit EZ DNA Methylation-Gold Kit (Zymoresearch, D5005). Bisulphite-converted DNA was used as a template for the targeted DNA amplification. PCR fragments were purified and used for sequencing. Adapters were removed from the deep sequenced PCR products by cutadapt tool (Martin, 2014). For alignment the reads, the target sequence was converted by Bismark aligner. Adapter trimmed sequences (90-100 bp) were aligned to target site using Bismark aligner tool with default parameters (Krueger and Andrews, 2011). DNA methylation status of the targeted site was extracted and coverage reports were generated using the Bismark aligner tool. The obtained results were analysed using methylation package ViewBS (Huang et al., 2018). Sequence data were deposited in the National Center for Biotechnology Information (NCBI) SRA database (Accession number: PRJNA1144360).

### Statistical analyses

Statistical analysis was performed using two-way analysis of variance (ANOVA) followed by the post hoc Tukey HSD test (P <0.05) using the Graphpad prism software (version 8). The data presented in graphs represent the mean value ± standard error of three independent experiments.

## Data availability statement

The data supporting the findings of this study are available upon request from the corresponding author, LS. The raw sequencing data have been deposited in the NCBI and can be accessed RNA-seq: https://dataview.ncbi.nlm.nih.gov/object/PRJNA1144338?reviewer=60o8cel20hm9nhfovce9oitou5. ChIP-seq: https://dataview.ncbi.nlm.nih.gov/object/PRJNA1144218?reviewer=qnlmhdp73uksjnvl3ju8jgv18e, and Targetted bisulphite sequencing: https://dataview.ncbi.nlm.nih.gov/object/PRJNA1144360?reviewer=t3v21vbt3fpdem9f12100qgsu.

## Author contributions

MP and LS conceived and designed the study and wrote the manuscript with contributions from all the authors. MP conducted the experiments and analyzed the data. MP, KK, LZ and GR helped with plant growth and physiological analysis. PS helped with the generation of RNA-seq data. AP performed DNA methylation analysis. IN performed NGS. LS supervised the experiments. MP, LS and PVS interpreted the results. All authors contributed to the article and approved the submitted version.

## Acknowledgments

Authors are indebted for Annamária Király for her assistance in optimization of rapeseed cultures.

## Supplementary material

**Figure S1:** Response of rapeseed to osmotic stress.

**Figure S2.** GO analysis of differentially regulated genes after 6 h of PEG treatment.

**Figure S3.** KEGG analysis of differentially regulated genes after 6h of PEG treatment.

**Figure S4.** GO analysis of differentially regulated genes after 24 h of PEG treatment.

**Figure S5.** KEGG analysis of differentially regulated genes after 24 h of PEG treatment.

**Figure S6.** GO analysis of differentially regulated genes after 6 h and 24 h of PEG treatment.

**Figure S7.** KEGG analysis of differentially regulated genes after 6 h and 24 h of PEG treatments.

**Figure S8:** Gene Ontology and KEGG analysis of genes Up-regulated 6 h and Down-regulated 24 h.

**Figure S9:** Heatmap of differentially expressed transcription factors.

**Figure S10:** Heatmap of differentially expressed chromatin remodelers.

**Figure S11**: GO analysis of genes differentially regulated at 6 and 24 hours of PEG treatment.

**Figure S12:** ChIP-seq analysis of rapeseed in response to PEG treatment.

**Figure S13.** GO analysis of differentially H3K4me3 enriched regions.

**Figure S14.** GO analysis of differentially H3K27me3 enriched regions.

**Figure S15.** Transposons showing differential enrichment of H3K4me3 marks.

**Figure S16.** Transposons showing differential enrichment of H3K27me3 marks.

**Figure S17:** IGV image showing ChIP-Seq and RNA-seq tracks of H3K4me3 and H3K27me3 enrichments

**Figure S18:** Heatmap of the RNAseq and ChIP-seq data of 58 selected genes.

**Figure S19:** Asymmetrical distribution of epigenomic marks in the An and Cn subgenomes of *Brassica napus*.

**Figure S20:** H3K4me3 and H3K27me3 enrichments of BnP5CSA and BnP5CSB genes.

**Table S1:** Primers for qRT-PCR analysis of stress responsive genes in *Brassica napus*

**Table S2**: Primers for ChIP-qRT-PCR analysis of BnP5CSA genes.

**Table S3**. RNA-seq quality metrics.

**Table S4**. ChIP-seq quality metrics.

**Table S5:** Primers for targetted bisulphite sequencing analysis of *BnP5CSA* genes.

**Table S6:** Transcript and histone modification data of proline metabolic genes.

## Supplementary Datasets

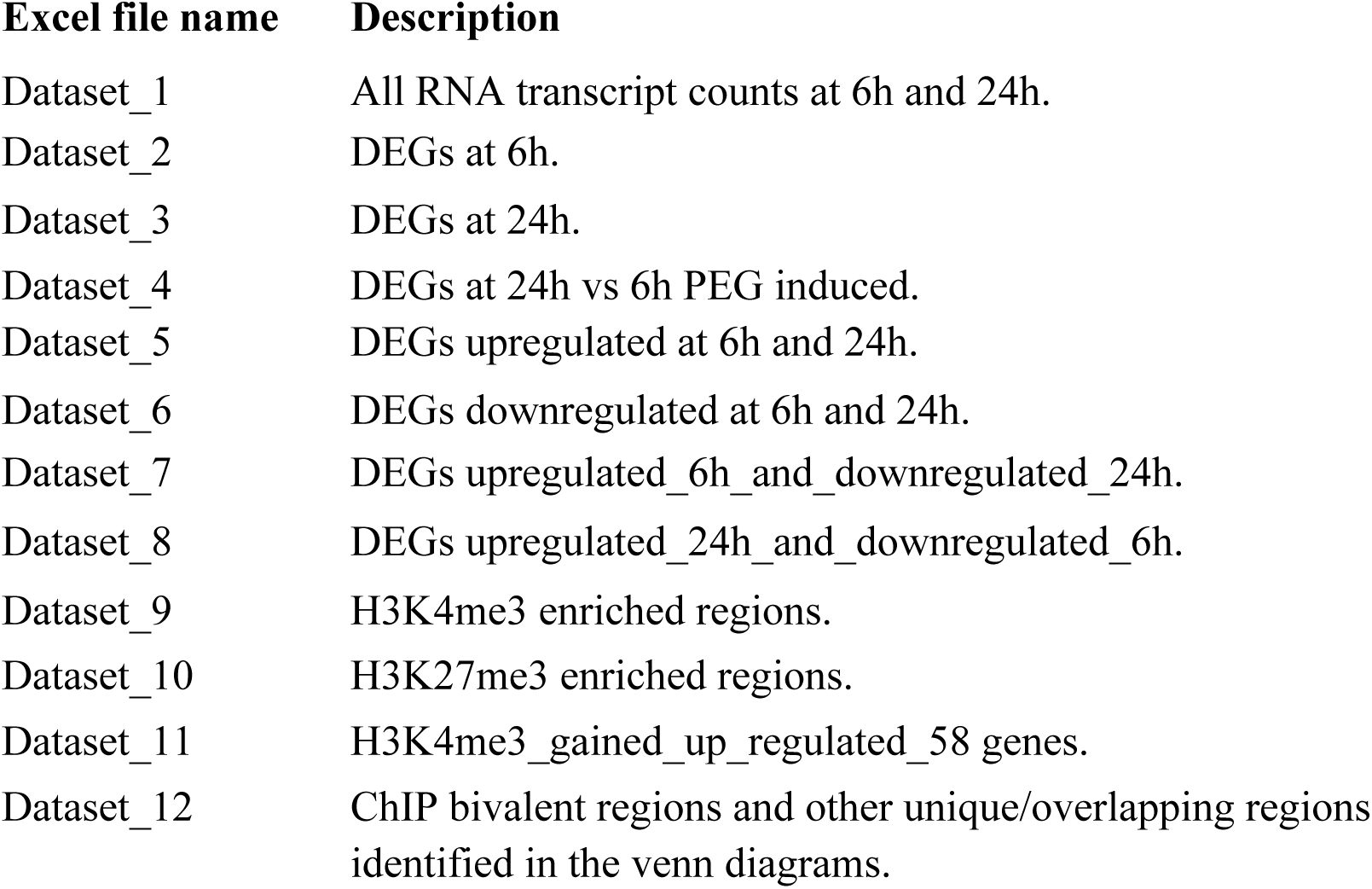

